# Influenza A virus during pregnancy disrupts maternal intestinal immunity and fetal cortical development in a dose- and time-dependent manner

**DOI:** 10.1101/2023.12.18.572222

**Authors:** Ashley M. Otero, Meghan G. Connolly, Rafael J. Gonzalez-Ricon, Selena S. Wang, Jacob M. Allen, Adrienne M. Antonson

**Author notes:** Corresponding Author Address: 1201 W Gregory Dr, Urbana, IL 61801.

## Abstract

Epidemiological studies link neurodevelopmental disorders (NDDs) with exposure to maternal viral infection in utero. It is hypothesized that the mechanism governing this link involves the activation of maternal intestinal T helper 17 (T_H_17) cells, which produce effector cytokine interleukin (IL)-17. While IL-17 is implicated as a major driver of fetal brain abnormalities, this inflammation-induced T_H_17 pathway has not been thoroughly examined in models of live viral infection during pregnancy. Influenza A virus (IAV) infection is consistently linked to offspring NDDs and can result in host intestinal dysregulation. Therefore, it is possible that intestinal T_H_17 cells and subsequent production of IL-17 could drive fetal brain abnormalities during gestational IAV infection. To test this, we inoculated pregnant mice with two infectious doses of IAV and evaluated peak innate and adaptive immune responses in the dam and fetus. While respiratory IAV infection led to dose-dependent maternal colonic shortening and microbial dysregulation, there was no elevation in intestinal T_H_17 cells nor IL-17. Fetal cortical abnormalities and global changes in fetal brain transcripts were observable in the high-dose IAV group, despite a lack of IL-17 signaling. Profiling fetal microglia and border-associated macrophages (BAMs) –potential cellular mediators of IAV-induced cortical abnormalities –revealed dose-dependent differences in the numbers of BAMs but not microglia. Overall, our data support the idea of an infection severity threshold for downstream maternal inflammation and fetal cortical abnormalities, confirming the use of live pathogens in NDD modeling to better evaluate the complete immune response and to improve translation to the clinic.

## Introduction

Influenza A virus (IAV) is a highly contagious respiratory pathogen that annually infects 5-10% of the global population [1]. Most individuals have mild symptoms; however, IAV infection during pregnancy poses a substantially increased risk of morbidity and mortality in both mother and infant [2–4] due to pregnancy-mediated changes to the maternal immune landscape [5]. Gestational IAV infection also has the potential to cause long-lasting negative health outcomes in the developing offspring. Epidemiological studies demonstrate that IAV infection during pregnancy increases the prevalence of offspring neurodevelopmental disorders (NDDs) like schizophrenia [6–8], bipolar disorder [9], and autism spectrum disorder (ASD) [10].

Early rodent models of prenatal exposure to IAV found that offspring developed neuropathology similar to that seen in ASD and schizophrenia [11–13]. Notably, the cause of these brain abnormalities was determined to be from the maternal anti-viral inflammatory response rather than vertical transmission of the virus from mother to fetus [14]. Pathogen mimetics, such as bacterial endotoxin lipopolysaccharide (LPS) or synthetic dsRNA polyinosinic-polycytidylic acid (poly I:C), are currently popular choices for maternal immune activation (MIA) modeling due to the controlled and predictable innate immune response elicited, which enables researchers to target specific fetal developmental periods. While these mimetic models recapitulate NDD-like behavioral and neuropathological offspring phenotypes [15,16], they fail to recreate the full spectrum of pathogen-induced pathological conditions. Unlike poly I:C, live viruses like IAV actively replicate within infected tissue and elicit a complex cascade of innate *and* adaptive immune responses [17–19]. Thus, certain characteristics of important downstream immune signaling cascades differ between poly I:C and live IAV. For instance, poly I:C-initiated MIA models implicate interleukin (IL)-17 as the major driver of fetal brain abnormalities [20–22], whereby activation of pre-existing maternal intestinal T helper (T_H_)-17 cells leads to increased production of IL-17. While IL-17-producing intestinal T_H_17 cells have also been identified as potential drivers of IAV-mediated intestinal injury [23], this occurs on a very different time scale (6-8 days following IAV inoculation versus 24-28 hours following poly I:C injection) and requires differentiation and propagation of naïve CD4^+^ T cells into pathogenic T_H_17 cells [23,24]. Understanding these model differences is critical for effectively delineating the etiologies of offspring neurodevelopmental disease during MIA and for improving translation to the clinic.

Still, the mechanism by which maternal intestinal immune cells might perturb the fetal brain remains to be fully elucidated. Recent work indicates that elevated maternal IL-17 leads directly to fetal brain abnormalities by binding receptors on neurons, as indicated by increased transcription of neuronal IL-17RA [21]. However, it is unclear how this proposed receptor-ligand binding leads to fetal cortical malformations following maternal poly I:C challenge. Furthermore, it is unclear whether a similar signaling cascade is activated during maternal IAV infection or if there are additional inflammatory signals at play. Mounting evidence indicates that fetal microglia and border-associated macrophages (BAMs) also respond to maternally derived inflammation [25,26] and could be contributing to neocortical developmental abnormalities [27]. Microglia are present in the embryonic brain at the onset of neurogenesis [28], which places them at the center of early neuronal support. One of their many roles during development includes shaping the neocortex by stimulating neural precursor cell proliferation [29] and by subsequent phagocytosis of excess neural precursor cells [30]. At least one study directly implicates fetal microglia in mediating interneuron deficits during MIA [27]. Less is known about the role of BAMs during MIA, although recent evidence suggests they play a significant role in propagating MIA-induced inflammatory signaling at the embryonic choroid plexus [26]. Critically, fetal microglia and BAMs have never been examined during IAV-induced maternal inflammation.

In this study, we used an established rodent model of gestational viral infection with mouse-adapted IAV [31]. By comparing moderate and high IAV challenge doses in pregnant dams, we demonstrate that a maternal infection severity threshold exists for the onset of fetal brain abnormalities. Furthermore, while high-dose IAV induced cortical abnormalities and increased the number of BAMs in the fetal brain, IAV infection failed to increase circulating levels of maternal IL-17A and did not induce a pathogenic intestinal T_H_17 cell phenotype. These findings indicate that gestational IAV infection leads to aberrant fetal cortical development via IL-17-independent mechanisms.

## Materials and Methods

### Animals

Singly housed male and pair-housed nulliparous female C57BL/6NTac mice obtained from Taconic Biosciences (Germantown, NY) at 9-to-10 weeks old were acclimated to the University of Illinois Urbana-Champaign animal facilities for a minimum of one week. Following acclimation, mice were trio-bred for 2-to-4 days. The presence of a vaginal plug was designated as gestational day (GD)0.5. Animals were maintained on a 12 h light–dark cycle, and body weights were recorded daily. A total of 60 pregnant dams across five biological replicates were used to examine tissues at 2 days post inoculation (dpi; GD11.5). 29 pregnant dams across three biological replicates were used to examine tissues at 7 dpi (GD16.5). A separate cohort of 20 pregnant dams was used for bulk RNA-sequencing of the fetal brain at 7 dpi. All animal research was approved by and performed in accordance with The Institutional Animal Care and Use Committee (IACUC) at the University of Illinois Urbana-Champaign.

### Influenza A viral inoculation

Mouse-adapted IAV (subtype H3N2, strain X31) was provided by Dr. Jacob Yount at The Ohio State University. On GD9.5, pregnant dams were randomly assigned to infected or control groups. Murine infection at GD9.5 approximates the end of the first trimester in humans [32], a time when the risk of IAV infection leading to aberrant offspring neurodevelopmental outcomes is high [7]. They were then anesthetized by inhalation of isoflurane before intranasal inoculation with either a moderate dose of 10^3^ tissue culture infectious dose (TCID_50_) IAV (X31_mod_) or a high dose of 10^4^ TCID_50_ IAV (X31_hi_) in sterile saline. Control animals were inoculated with sterile saline (Con).

### Tissue collection

Tissues were collected and examined at either 2 or 7 dpi. Pregnant dams were euthanized by CO_2_ inhalation, and tissues were excised under sterile conditions. Maternal whole blood was obtained through blind cardiac puncture and subsequently centrifuged at 2,000 x g at 4°C for 10 min. Maternal serum was aliquoted and stored at -80°C until analysis. Dams whose tissues were used for immunohistochemistry at 2 dpi were transcardially perfused with 25 mL sterile saline following blood collection. The uterus was removed from the dam and placed in ice cold phosphate-buffered saline (pH = 7.4; PBS), then placentae and fetuses were separated and cleaned. Fetuses were immersion fixed in 10% neutral buffered formalin (NBF) and stored at 4°C. The ileum, spleen, and right lung were collected from each pregnant dam and snap frozen on dry ice before storing at -80°C until further processing. The left lung was immersion fixed in 10% NBF and stored at 4°C until histopathology processing. Maternal colon length was measured before collection. The colon was then snap frozen and stored at -80°C or placed in ice cold media for further processing (see flow cytometry section).

### Lung Histology

Dam lung tissue was processed at the University of Illinois Urbana-Champaign Veterinary Diagnostic Laboratory. Tissues were paraffin-embedded and stained with hematoxylin and eosin (H&E). Slides were evaluated by a board-certified veterinary comparative pathologist, Dr. Shih-Hsuan Hsiao, who was blinded to treatment groups. Semi-quantitative scoring of lung histopathology was performed using a rubric from previously published scoring methods [33]. The following lesions were independently scored for each sample: bronchitis, interstitial inflammation, edema, endothelialitis, pleuritis, and thrombus formation. Each lesion type was graded on a scale of 0 to 4 (0: absent, 1: mild, 2: moderate, 3: severe, 4: very severe). The total histopathological score is expressed as the sum of scores for all parameters with a maximum score of 24.

### Quantitative real-time PCR (qPCR)

RNA was isolated using TRIzol Reagent per the manufacturer’s protocol (Invitrogen, Carlsbad, CA, Catalog no. 15-596-026). RNA integrity and concentration were determined using the Nanodrop Nd-8000 Spectrophotometer (Thermo Fisher Scientific). Four µg of cDNA per sample was synthesized using the High-Capacity cDNA Reverse Transcription Kit (Applied Biosystems, Foster City, CA, Catalog no. 43-688-13) and Mastercycler Pro Thermal Cycler (Eppendorf) and was subsequently diluted 1:5 in DEPC water. qPCR was performed on a QuantStudio 5 Real-Time PCR System (Thermo Fisher Scientific) using PowerTrack SYBR Green Master Mix (Applied Biosystems, Catalog no. A46111). Data were analyzed using the 2^-ΔΔCt^ method against housekeeping genes (for mouse cDNA) or total Eubacteria (for microbial DNA) and presented as relative expression compared to control. Custom primer information is listed in **Supp. Table S1** (Integrated DNA Technologies, Coralville, IA). Statistics were run on relative expression values.

### Protein immunoassays

Protein was isolated using Tissue Protein Extraction Reagent (T-PER) per the manufacturer’s protocol (Thermo Fisher Scientific, Waltham, MA, Catalog no. 78510). Protein concentration was measured using Pierce BCA Protein Assay (Thermo Fisher Scientific, Catalog no. PI23225) and read on BioTek ELx800 Microplate Reader (Agilent). Serum samples were analyzed using LEGENDplex MU Inflammatory Panel (13-plex) per the manufacturer’s protocol (BioLegend, San Diego, CA, Catalog no. 740446). All samples were run on the Attune NxT Flow Cytometer (Thermo Fisher Scientific) and analyzed on Qognit software (San Carlos, CA).

### Immunohistochemistry

Upon tissue collection, embryonic day (E)11.5 fetuses and E16.5 fetal heads were immersion-fixed in 10% NBF for 24 h at 4°C. They were then decanted and washed twice in PBS followed by immersion in 30% sucrose solution with sodium azide for at least 48 h at 4°C. E16.5 brains were dissected from heads after fixation and cryoprotection; E11.5 fetuses were kept whole. Cryoprotected tissue was embedded in OCT tissue-tek (Thermo Fisher Scientific, Catalog no. NC9159334) and stored long-term at -80°C. 60µm sagittal sections of E11.5 fetuses and 25µm coronal sections of E16.5 fetal brains were cryosectioned. Free-floating batch staining was performed for each staining configuration. Sections were washed three times in PBS with 0.05% Tween-20 (Thermo Fisher Scientific, Catalog no. PRH5152; PBST) for 5 min each time and subsequently incubated with blocking buffer (5% goat serum [R&D Systems, Minneapolis, MN, Catalog no. S13110], 1% bovine serum albumin [Thermo Fisher Scientific, Catalog no. 126609100GM], 0.3% Triton-X 100 [Thermo Fisher Scientific, Catalog no. ICN19485450] in PBST) for 1 h at room temperature. For stains with a mouse host, sections were blocked with Mouse-on-Mouse IgG Blocking Solution for 1 h at room temperature (1:30; Thermo Fisher Scientific, Catalog no. R37621). Sections were incubated overnight at 4°C with primary antibodies—rabbit anti-Iba1 (1:1000; Wako Chemicals U.S.A, Richmond, VA, Catalog no. 019-19741), rat anti-CD206 (1:1000; Biorad, Hercules, CA, Catalog no. MCA2235GA), rat anti-Ki67 (1:500; Invitrogen, Catalog no. 14-5698-82), rabbit anti-TBR1 (1:1000; Abcam, Cambridge, UK, Catalog no. ab183032), or mouse anti-SATB2 (1:300; Abcam, Catalog no. ab51502). Sections were washed three times in PBST and incubated with secondary antibodies—Alexa Fluor 594 goat anti-rabbit IgG H&L (1:250; Jackson ImmunoResearch, West Grove, PA, Catalog no. 111-585-003), Alexa Fluor 488 goat anti-rat IgG H&L (1:250; Jackson ImmunoResearch, Catalog no. 112-545-003), or Alexa Fluor 488 goat anti-mouse IgG H&L (1:400; Thermo Fisher Scientific, Catalog no. A-11001)—for 2 h followed by staining in DAPI (Thermo Fisher Scientific, Catalog no. EN62248) for 1 min. Sections were mounted with Fluoromount-G Mounting Medium (Thermo Fisher Scientific, Catalog no. 5018788) and stored long-term at 4°C.

### Imaging & Image Analysis

All images were acquired using a ZEISS AxioScan.Z1 slide scanner with system configurations as follows: Colibri 7 LED light source: 385 nm, 430 nm, 511 nm, 555 nm, 590 nm, 630 nm; Objectives: 5x/0.25 10x/0.45, 20x/0.8, 40x/0.5 Pol and 50x/0.8 Pol; Filter: GFP, DsRed, Cy5, DAPI/GFP/CY3/Cy5, and CFP/FP/mCherry; Camera: Hamamatsu Orca Flash, AxioCam IC (CCD AxioCam IC Color camera) and Hitachi HV-F202SCL. Images were subjected to the same conditions for each staining configuration. Image files were blinded and analyzed using ZEISS ZEN 3.0 Blue software (Oberkochen, DE). Cell counts were performed manually and normalized by area (mm^2^) for Iba1^+^CD206^-^ (microglia), Iba1^+^CD06^+^ (BAMs), and Iba1^+^Ki67^+^ (proliferating Iba1^+^ cells) in E11.5 and 16.5 fetal brains. Mean fluorescence intensity (MFI) was performed using ImageJ on a 300 x 300µm^2^ region of interest (ROI) for SATB2 (upper excitatory neurons) and TBR1 (deep excitatory neurons) in E16.5 fetal brains. The ROI was further divided into 10 equal laminar bins, and the signal intensity of each bin was normalized relative to the total signal intensity of the ROI. The location of the primary somatosensory cortex region was determined based on the distance from the retrosplenial cortex relative to the length of the dorsal midline. E11.5 brain regions were identified using Chen et al. (2017) [34] and Kaufman’s Atlas of Mouse Development [35]. E16.5 brain regions were identified using Uta Schambra’s Prenatal Mouse Brain Atlas [36].

### Flow Cytometry

Flow cytometry was performed on 2 and 7 dpi colonic lamina propria lymphocytes (LPLs) based on previously published methods [37]. The colon was flushed with FACS buffer (2% FBS [Thermo Fisher Scientific, Catalog no. MT35010CV], 2mM EDTA [Thermo Fisher Scientific, Catalog no. 15-575-020] in 1xHBSS [Thermo Fisher Scientific, Catalog no. 14175145]). Mesenteric fat was removed, and the colon was longitudinally bisected. Colons were incubated in EDTA-DTT buffer (2% FBS, 1mM EDTA, 1mM DTT [Sigma Aldrich, St. Louis, MO, Catalog no. 10708984001] in 1xHBSS) at 37°C, 250 rpm for 20 min to remove epithelial cells and intraepithelial lymphocytes. Colons were subsequently incubated in digestion buffer (2% FBS, 50µg/mL DNase I [Sigma Aldrich, Catalog no. 10104159001], and 62.5µg/mL Liberase [Sigma Aldrich, Catalog no. 5401127011] in RPMI-1640 [Corning, Corning, NY, Catalog no. 15-040-CV]) at 37°C, 250 rpm for 45 min. The resulting cells were isolated using a 100µm cell strainer followed by a 40/80 Percoll (Thermo Fisher Scientific, Catalog no. 45001753) gradient. Cells were resuspended in T cell culture media (10% FBS, 50µg/mL gentamicin sulfate [Corning, Catalog no. 30-005-CR], 2x GlutaMAX [Thermo Fisher Scientific, Catalog no. 35050061], 1xPenicillin/Streptomycin [Corning, Catalog no. 30002Cl], and 55µM 2-Beta-Mercaptoethanol [Thermo Fisher Scientific Catalog no. 21-985-023] in RPMI-1640) and counted using Invitrogen Countess Automated Cell Counter and adjusted so there were 1×10^6^-2×10^6^ cells/100µl. Cells were then cultured for 3 h in T cell stimulation media (1000ng/mL PMA [Sigma Aldrich, Catalog no. P1585-1MG], 2µM Ionomycin [Sigma Aldrich, Catalog no. I0634-1MG], and 2µg/mL GolgiPlug [BD Biosciences, Franklin Lakes, NJ, Catalog no. 555029] in T cell culture media) at 37°C, 5% CO_2_. Cells were stained with LIVE/DEAD Fixable Aqua (Thermo Fisher Scientific, Catalog no. L34966), Rat anti-mouse CD16/32 Fc Block (BD Biosciences, Catalog no. 553142), and surface markers CD4 anti-mouse BB700 (BD Biosciences, Catalog no. 566407) and CD45 anti-mouse APC-Cy7 (BD Biosciences, Catalog no. 557659). Cells were then fixed and permeabilized using FOXP3 Transcription Factor Staining Buffer Set (Thermo Fisher Scientific, Catalog no. 00-5523-00) and stained for intracellular markers RORγt anti-mouse PE-CF594 (BD Biosciences, Catalog no. 562684), Tbet anti-mouse APC (Thermo Fisher Scientific, Catalog no. 17-5825-82), IL-17A anti-mouse Alexa Fluor 488 (BioLegend, Catalog no. 506910), IL-17F anti-mouse PE (BD Biosciences, Catalog no. 561627), and IFNγ anti-mouse Brilliant Violet 421 (BioLegend, Catalog no. 505830). Cells were run on the Attune NxT Flow Cytometer (Invitrogen). Fluorescence minus ones (FMOs) were used daily for gating. Compensation was done using UltraComp compensation beads (Thermo Fisher Scientific, Catalog no. 501129040). Flow analysis was done using FlowJo (Ashland, OR).

### 16S Sequencing

DNA extraction was performed using a QIAamp Fast DNA Stool Mini Kit (Qiagen, Valencia, CA, Catalog no. 51604) following manufacturer’s instructions, with slight modifications as previously described [38]. Briefly, 20-40 mg of stool was incubated for 45 min at 37°C in lysozyme buffer (22 mg/ml lysozyme, 20 mM TrisHCl, 2 mM EDTA, 1.2% Triton-x, pH 8.0), then bead-beat for 150 s with 0.1 mm zirconia beads. Samples were incubated at 95°C for 5 min with InhibitEX Buffer, then incubated at 70°C for 10 min with Proteinase K and Buffer AL. Following this step, the QIAamp Fast DNA Stool Mini Kit isolation protocol was followed, beginning with the ethanol step. DNA was quantified with the Qubit 2.0 Fluorometer (Life Technologies, Carlsbad, CA) using the dsDNA Broad Range Assay Kit.

After extraction and DNA quality assurance through gel electrophoresis, library construction was completed using a Fluidigm Access Array system in the Functional Genomics Unit of the Roy J. Carver Biotechnology Center at the University of Illinois Urbana-Champaign. After library construction, 250 bp of the V4 region of the 16SrRNA gene were amplified and sequenced at the WM Keck Center for Biotechnology at the University of Illinois Urbana-Champaign using an Illumina MiSeq2000. The V4 region of the 16S rRNA gene was amplified using primers 515F (5’-GTGYCAGCMGCCGCGGTAA-3’) and 806R (5’-GGACTACNVGGGTWTCTAAT-3’). PCR reactions were conducted in triplicate and resulting amplicons were pooled.

Illumina libraries were generated from the pooled amplicons and paired-end (2 x 250 nt) sequencing was performed on an Illumina MiSeq. After sequencing and barcode trimming, raw sequence data (FASTQs) underwent quality control using DADA2, trimming low-quality bases with a cutoff Phred score >30. DADA2 was used for denoising, merging paired-end reads, and inferring amplicon sequence variants (ASVs). ASVs were mapped to SILVA rRNA database version 138.1 with QIIME2. Alpha diversity metrics (Shannon’s Index) were calculated to assess within-sample diversity. Beta diversity metrics (Unweighted Unifrac) were calculated to assess between-sample diversity. Downstream statistical analysis and data visualization were completed with MicrobiomeAnalyst. Taxa abundance data were filtered as follows: minimum count = 4, prevalence in samples = 20%, and percentage to remove = 10%. Data were then transformed using the centered log-ratio (CLR) method to mitigate compositional biases and improve interpretability. Negative binomial regression models were employed to explore relationships between dam influenza status and the gut microbiome.

### Bulk RNA-Sequencing

RNA from pooled fetal brains was extracted using the RNeasy Mini Kit (Qiagen, Catalog no. 74004). RNA quality and integrity were then determined by 28S/18S rRNA analysis with the Agilent 2100 Bioanalyzer. All samples scored an RNA Integrity Number over 7 indicating little to no signs of degradation. RNA-sequencing was performed at the Roy J. Carver Biotechnology Center. RNA-Seq libraries were prepared using the KAPA Stranded RNA-Seq Library Preparation Kit with an average fragment length of 100 bp. Libraries were pooled and quantified using qPCR and were then sequenced on one S Prime (SP) lane for 101 cycles from one end of the fragments on a NovaSeq 6000 (Illumina). FASTQ files were generated from the raw sequencing runs and demultiplexed with the bcl2fastq v2.20 Conversion Software (Illumina). Adaptors were trimmed from the 3’-end of the reads, and QC indicated no adaptor sequence contamination.

All reference files were downloaded from the National Center for Biotechnology Information (NCBI) ftp site. The *Mus musculus* H3N2 transcriptome file “GCF_000001635.27_GRCm39_rna.fna.gz; influenza_A_X-31_H3N2_rna.fa” from NCBI Annotation 109 was used for quasi-mapping and count generation. This transcriptome is derived from genome GRCm39 (Mouse); A/X-31(H3N2). Since the quasi-mapping step only uses transcript sequences, the gene model file “GCF_000001635.27_GRCm39_genomic.gff.gz; influenza_A_X-31_H3N2.gff3” was used to generate transcript-gene mapping table (“trx_EGids_GRCm39_annot109.txt”) for gene-level counts.

Salmon version 1.4.0 was used to quasi-map reads to the transcriptome and quantify the abundance of each transcript. The transcriptome was first indexed using the decoy-aware method in Salmon with the entire genome file “GCF_000001635.27_GRCm39_genomic.fna.gz; influenza_A_X-31_H3N2_genes.fa” as the decoy sequence. Then quasi-mapping was performed to map reads to the transcriptome with additional arguments --seqBias and --gcBias to correct sequence-specific and GC content biases, --numBootstraps=30 to compute bootstrap transcript abundance estimates, and --validateMappings to help improve the accuracy of mappings. Gene-level counts were then estimated based on transcript-level counts using the bias-corrected counts without an offset method from the ‘tximport’ package in R. The percentage of reads mapped to the transcriptome for all samples was between 70-80% (**Fig. S1A**). Normalization of samples was between 0.97-1.03 using the trimmed mean of M values (TMM) method (**Fig. S1B**). Remove Unwanted Variation (RUV) analysis was performed on the data. RUV is a method to estimate factors that can be added to the statistical model (co-variates) assuming that these factors are spurious technical variations in the samples and not biologically related. Additional normalization using RUV is often necessary for RNA-Seq experiments and helps to improve biological insights [39]. Clustering after RUV removal was done using the limma package in R (**Fig. S1C**). The limma-trend method was used to find differentially expressed genes using a model of ∼ treatment + 4 RUV factors [40]. P-values of differentially expressed genes were adjusted using Benjamini-Hochberg’s correction for false discovery (p<0.1). The Database for Annotation, Visualization, and Integrated Discovery (DAVID) was used to find the top ten significantly enriched up and downregulated Gene Ontology (GO) pathways.

### Statistics

Dam was treated as the experimental unit for all outcomes, and one representative fetus per litter was used for each experimental outcome. All data were analyzed using GraphPad Prism 9 Software (San Diego, CA) with significance set at alpha = 0.05 unless otherwise specified. Dam body weights across time were analyzed using repeated measures two-way ANOVA with Geisser-Greenhouse correction for sphericity and Tukey correction for multiple comparisons. Con, X31_mod_, and X31_hi_ groups were compared using one-way ANOVA assuming Gaussian distribution and homogeneity of variance of residuals, and Tukey correction for multiple comparisons was used. For data with unequal variance, Brown-Forsythe ANOVA was performed with Dunnet T3 correction for multiple comparisons. For non-parametric data, Kruskal-Wallis ANOVA was performed with Dunn’s correction for multiple comparisons. Kruskal-Wallis was used in the case that residuals did not meet normality or homogeneity of variance. Outliers were identified and removed using the ROUT method with Q=1%.

## Results

### Respiratory IAV infection leads to maternal inflammation at the site of infection and systemically in a dose- and time-dependent manner

To capture the complete immune response, we evaluated pregnant dams and fetuses at 2 and 7 dpi to approximate the peak of both innate and adaptive anti-viral responses, respectively [41,42]. We previously reported that fetal brain neuroinflammatory transcripts are unaltered by inoculation with a moderate infectious titer of IAV (X31_mod_) [31]; thus, we included an additional high infectious titer (X31_hi_) in the current study (**Fig. 1A**). To determine whether the infectious dose of IAV impacts maternal symptomology and pathology, dam body weights were monitored daily (**Fig. S2**), and anti-viral immune response was examined in the lungs and systemically. Dam body weight gain per day was significantly decreased from 2-to-7 dpi in X31_hi_ dams whereas X31_mod_ dams exhibited a decrease in body weight gain per day only at 6 dpi (**Fig. 1B**). Both moderate- and high-dose dams demonstrated a spike in body weight gain per day at 5 dpi followed by a subsequent dip at 6 dpi, which could indicate a potential shift from innate to adaptive immunity [42]. IAV infection did not affect litter size or number of fetal resorptions at either time point, which is consistent with our previous findings [31] (**Supp. Table S2-S3**). The presence of viral RNA was confirmed in infected maternal lungs, although there was no difference in gene expression between high and moderate doses (**Fig. 1C**). Histopathological scoring of H&E-stained maternal lungs showed increased total lung lesion scores regardless of viral dose (**Fig. 1D-E**). Specifically, scores for bronchitis, interstitial inflammation, and endothelialitis were upregulated throughout the infection (**Supp. Table S4-S5**). Genes encoding for classic pro-inflammatory cytokines IL-6, IL-1β, and tumor necrosis factor-alpha (TNF-α) were upregulated in the lungs in a dose-dependent manner at 2 dpi (**Fig. 1F**). Only *Il1b* remained elevated at 7 dpi (**Fig. 1G**). Type II antiviral interferon-gamma (IFNγ) was upregulated in X31_hi_ dams only during acute infection (**Fig. S3A**). Type I and type II interferons were downregulated in IAV-infected lungs at 7 dpi, which is consistent with prior kinetic studies of IAV-X31 [43] (**Fig. S3B**). We then evaluated IL-17 expression in the lungs. Elevated IL-17 signaling during IAV infection is shown to cause acute lung injury while also playing a protective role against secondary bacterial infections [44,45]. In line with these findings, we observed an increase in *Il17f* at 2 dpi (**Fig. 1H**) and an increase in *Il17a* and *Il17f* at 7 dpi in a dose-dependent manner (**Fig. 1I**).

**Figure 1.**
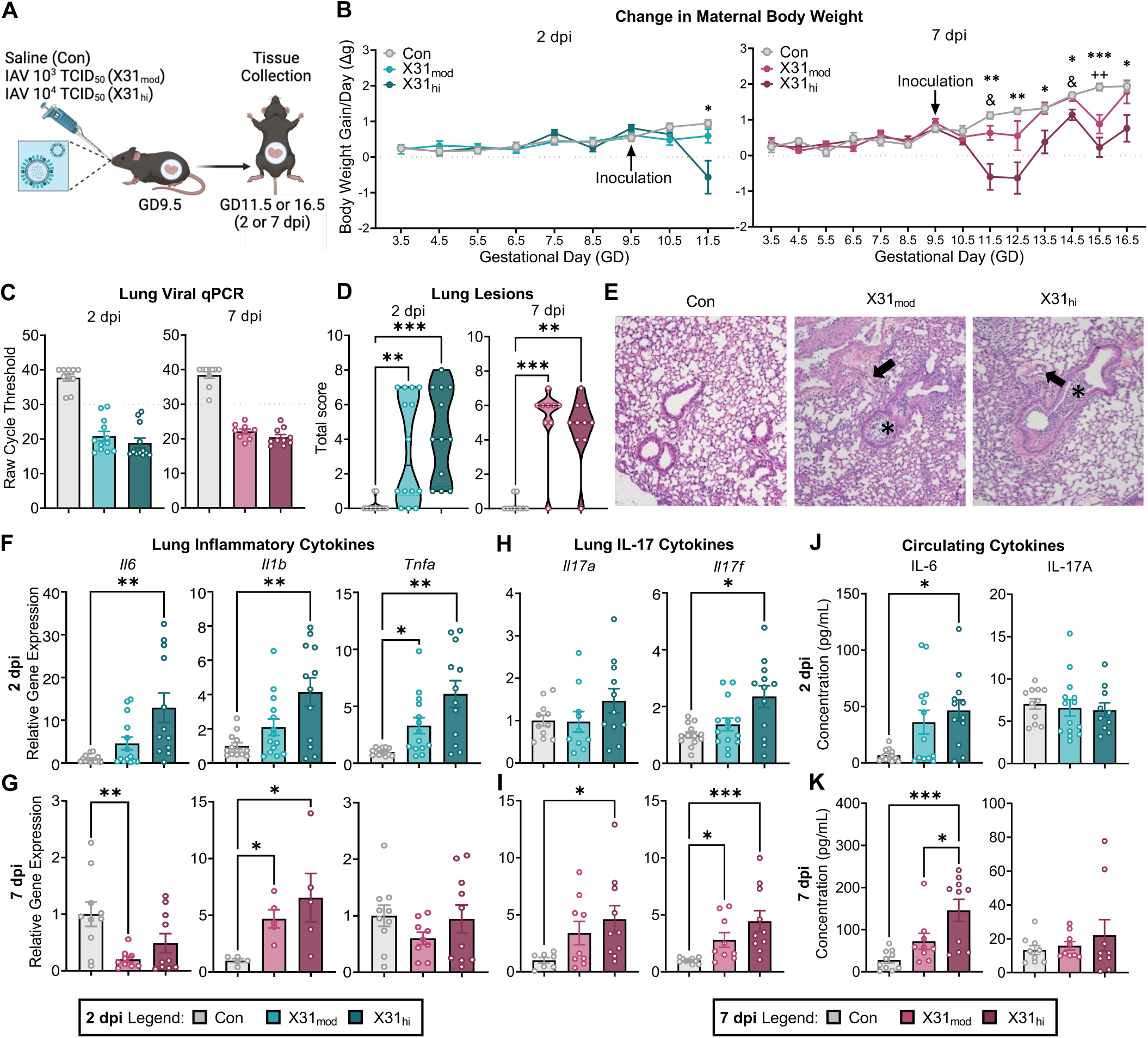
Respiratory IAV infection alters maternal lung inflammation and circulating cytokines in a dose- and time-dependent manner. **(A)** Experimental schematic. **(B)** IAV inoculation at GD9.5 suppressed body weight gain per day from 2-to-7 dpi in X31_hi_ dams only (repeated measures 2-way ANOVA, the main effect of time = p < 0.001; * = Con vs X31_hi_, & = X31_mod_ vs X31_hi_, + = Con vs X31_mod_). **(C)** The presence of IAV-X31 in lungs at 2 and 7 dpi was evaluated using qPCR with a cycle threshold of ≤ 30 cycles as confirmed infection (dotted line). **(D)** Quantification of H&E pathological scoring showed elevated lung lesion scores in infected dams. The scoring criteria are listed in the methods with additional scoring values in Supp. Table S4-S5. **(E)** Representative photomicrographs of H&E-stained lung sections. Asterisks (*) indicate bronchi filled with clusters of neutrophils with cellular debris, and arrows (→) indicate arterial and venous endothelia with rolling neutrophils. **(F-G)** Genes encoding for classic pro-inflammatory cytokines IL-6, IL-1β, and TNF-α in maternal lungs were **(F)** upregulated in a dose-dependent manner at 2 dpi whereas **(G)** only *Il1b* was upregulated at 7 dpi. **(H-I)** IL-17 genes were upregulated in the maternal lung at **(H)** 2 and **(I)** 7 dpi in a dose-dependent manner. **(J-K)** Maternal cytokines in circulation at **(J)** 2 and **(K)** 7 dpi. Pro-inflammatory cytokine IL-6 was upregulated in moderate- and high-dose dams proportional to dosage at both time points. IL-17A was not upregulated in circulation at either endpoint. *IAV* = influenza A virus, *GD* = gestational day, *dpi* = days post-inoculation, *Con* = saline control, *X31_mod_* = IAV-X31 10^3^ TCID_50_, *X31_hi_* = IAV-X31 10^4^ TCID_50_. Groups were compared using one-way ANOVA with Tukey post hoc for multiple comparisons. For data containing residuals with unequal variance, Brown-Forsythe and Welch’s ANOVA with Dunnett T3 post hoc multiple comparisons was used. For non-parametric data, Kruskal-Wallis ANOVA with Dunn’s correction for multiple comparisons was used. Data are means ± SEM; one symbol = p < 0.05, two symbols = p < 0.01, three symbols = p < 0.001; dots represent individual dams; n = 9-14 per treatment group.

To evaluate the systemic impacts of IAV across viral replication, we measured maternal serum cytokine concentrations. IL-6 was upregulated in a dose-dependent manner at 2 and 7 dpi (**Fig. 1J-K**). Anti-inflammatory cytokine IL-10 differed across groups at 2 dpi with a trending increase at 7 dpi (**Fig. S3C-D**). IFNγ and IL-23–a cytokine required for the commitment and propagation of T_H_17 cells [46]–were also upregulated in a dose-dependent manner at 7 dpi (**Fig. S3D**). Notably, IL-17A was not upregulated at either time point (**Fig. 1J-K**), which differs from poly I:C-induced MIA studies showing an elevation of IL-17A in maternal circulation [21,22]. However, this is consistent with a lack of elevated IL-17A in the serum of IAV-infected male mice [23].

### Respiratory IAV infection disrupts maternal intestinal immunity in a dose- and time-dependent manner

We evaluated the maternal intestines to determine if alterations in T_H_17 cells are present during gestational IAV, similar to poly I:C-induced MIA [21,22], and if these T_H_17 cells transition to a pathogenic phenotype, similar to IAV-infected male mice [23]. At 2 dpi, high-dose dams exhibit colonic shortening, which persists out to 7 dpi, a finding that has been found in previous mouse models of IAV [23,31] (**Fig. 2A**). Colonic shortening is a hallmark of intestinal inflammation and colitis [47]. Furthermore, while fewer T_H_17 cells reside in the colon compared to the small intestine, colonic T_H_17 cells are more susceptible to developing pathogenic phenotypes [24,48]. Therefore, we decided to phenotype colonic lamina propria T_H_17 cells in IAV-infected dams (**Fig. 2B**). CD45^+^CD4^+^ lymphocytes were gated on markers for homeostatic T_H_17 (RORγt, IL-17A, IL-17F), T_H_1 (Tbet, IFNγ), and pathogenic T_H_17 (RORγt, Tbet, IFNγ, IL-17A) cells [24,49]. We observed a downregulation in IL-17F^+^ and IL-17F^+^RORγt^+^ colonic T cells from X31_hi_ dams at 2 dpi (**Fig. 2C**). This finding persisted into 7 dpi with an additional decrease in RORγt^+^ and IL-17A^+^RORγt^+^ T cells (**Fig. 2D, S4A**). Further gating on CD45^+^CD4^+^RORγt^+^ and CD45^+^CD4^+^Tbet^+^ T cells revealed downregulation in classic T_H_17 (IL-17F^+^) cells at 2 and 7 dpi and no changes in classic T_H_1 cells, respectively (**Fig. S4B-E**). Additionally, there was no evidence to indicate an increase in pathogenic colonic T_H_17 cells when gating for RORγt^+^Tbet^+^ and IL-17A^+^IFNγ^+^ double-positive cells (**Fig. S4F-G**).

**Figure 2.**
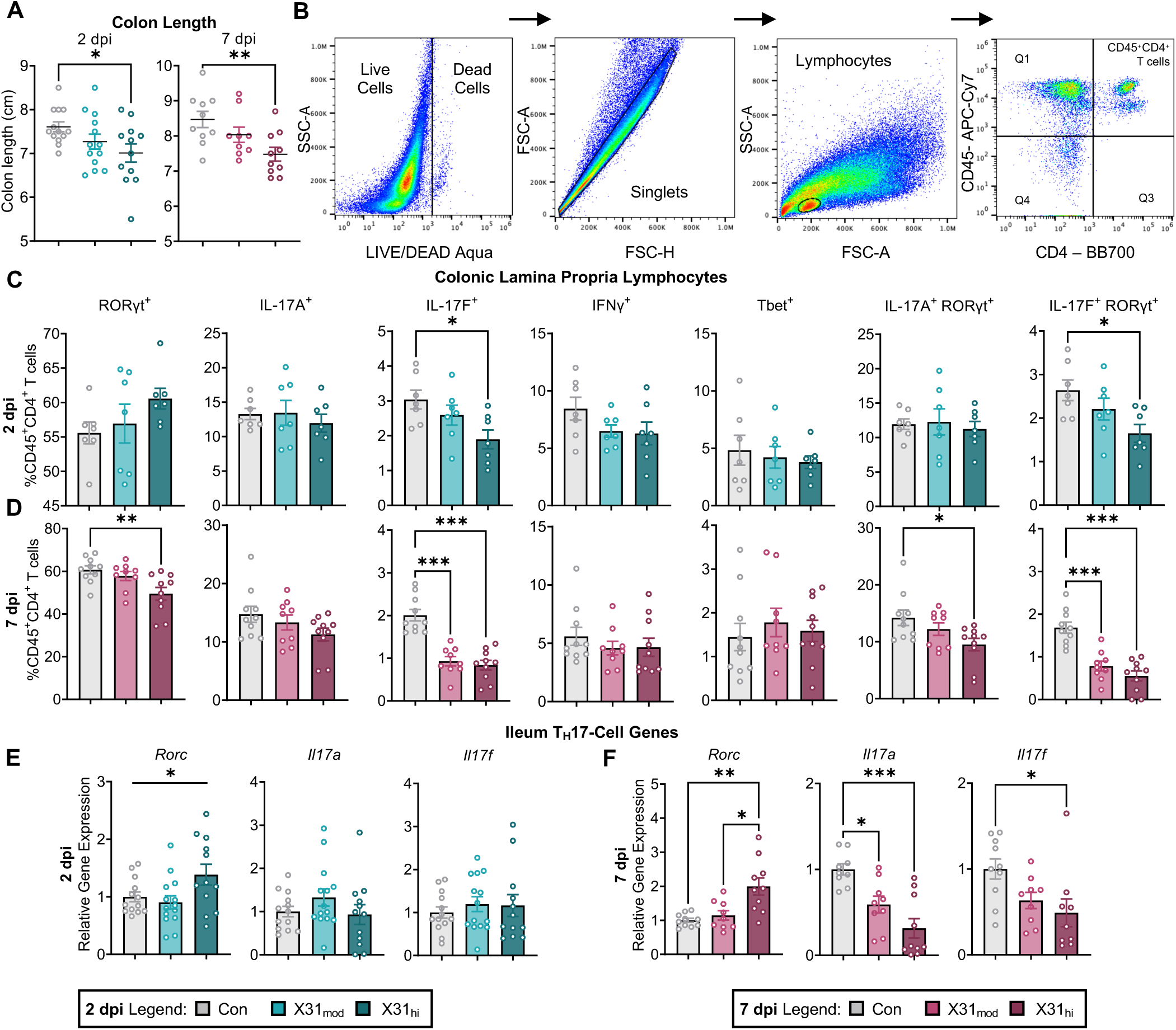
Respiratory IAV infection dysregulates maternal intestinal immunity in a dose- and time-dependent manner. **(A)** IAV reduced colon length in high-dose dams as early as 2 dpi, and this persisted at 7 dpi. **(B)** Example flow cytometry gating for colonic LPLs. **(C-D)** Gating on CD45^+^CD4^+^ T cells in the colon revealed **(C)** downregulation of IL-17F^+^ and IL-17F^+^RORγt^+^ T cells at 2 dpi which persisted into **(D)** 7 dpi in addition to downregulation of RORγt^+^ and IL-17A^+^RORγt^+^ T cells. **(E-F)** qPCR of the ileum, the terminal end of the small intestine, confirms findings in the colon where **(E)** little changes were observed at 2 dpi and **(F)** downregulation in *Il17a* and *Il17f* was seen at 7 dpi. Notably, *Rorc* transcription did not coincide with decreased RORγt protein expression. *IAV* = influenza A virus, *dpi* = days post-inoculation, *Con* = saline control, *X31_mod_* = IAV-X31 10^3^ TCID_50_, *X31_hi_* = IAV-X31 10^4^ TCID_50,_ *SSC-A* = side scatter-area, *FSC-A* = forward scatter-area, *LPL* = lamina propria lymphocytes. Groups were compared with one-way ANOVA with Tukey post hoc for multiple comparisons. For data containing residuals with unequal variance, Brown-Forsythe and Welch’s ANOVA with Dunnett T3 post hoc multiple comparisons was used. For non-parametric data, Kruskal-Wallis ANOVA with Dunn’s correction for multiple comparisons was used. Data are means ± SEM; * = p < 0.05, ** = p < 0.01, *** = p < 0.001; dots represent individual dams; n = 7-14 per treatment group.

To determine if the downregulation of IL-17 was specific to the colon, we looked at gene expression in the ileum of the small intestine, which is where the majority of homeostatic T_H_17 cells reside [50,51] and the typical intestinal region of interest in MIA studies [20,21]. qPCR of the ileum revealed a similar pattern to colonic flow cytometry findings: no transcripts differed at 2 dpi whereas both *Il17a* and *Il17f* decreased in the IAV groups at 7 dpi (**Fig. 2E-F**). There was also a decrease in ileal *Il22*, another cytokine produced by T_H_17 cells (**Supp. Table S7**). Contrastingly, transcription of *Rorc*, the gene encoding for T_H_17 cell transcription factor RORγt, was upregulated at both time points in high-dose dams only (**Fig. 2E-F**). We previously observed this same upregulation in *Rorc* in both the colon and ileum of moderate-dose dams at 7 dpi [31]. Thus, increased transcription of *Rorc* does not coincide with an increased number of RORγt^+^ cells in this model. We then looked upstream at T_H_17-cell priming genes and found upregulation in ileal *Il6*, downregulation in *Tgfb1* and *Il1b*, with no changes in *Il23a* at 7 dpi (**Supp. Table S7**). Altogether, these data demonstrate that gestational IAV alters intestinal immunity throughout infection despite a lack of viral replication within the intestines (**Supp. Table S6-7**). Notably, it appears that maternal colonic shortening cannot be explained by IL-17-producing T_H_17 cells in our model of IAV infection.

### Respiratory IAV infection alters the maternal colonic microbiome

Intestinal T_H_17 cells rely on segmented filamentous bacteria (SFB), which regulate the development of homeostatic T_H_17 cell populations in the small intestine [51]. The presence of SFB in pregnant mice is necessary to induce activation of T_H_17 cells and is also required for neuropathological and behavioral abnormalities in offspring from poly-I:C-induced MIA [20,22]. Contrastingly, we observed a decrease in SFB gene expression in X31_hi_ dams at 7 dpi (**Fig. 3A**). This corroborates what was previously found in the intestinal contents of IAV-infected male mice [23].

**Figure 3.**
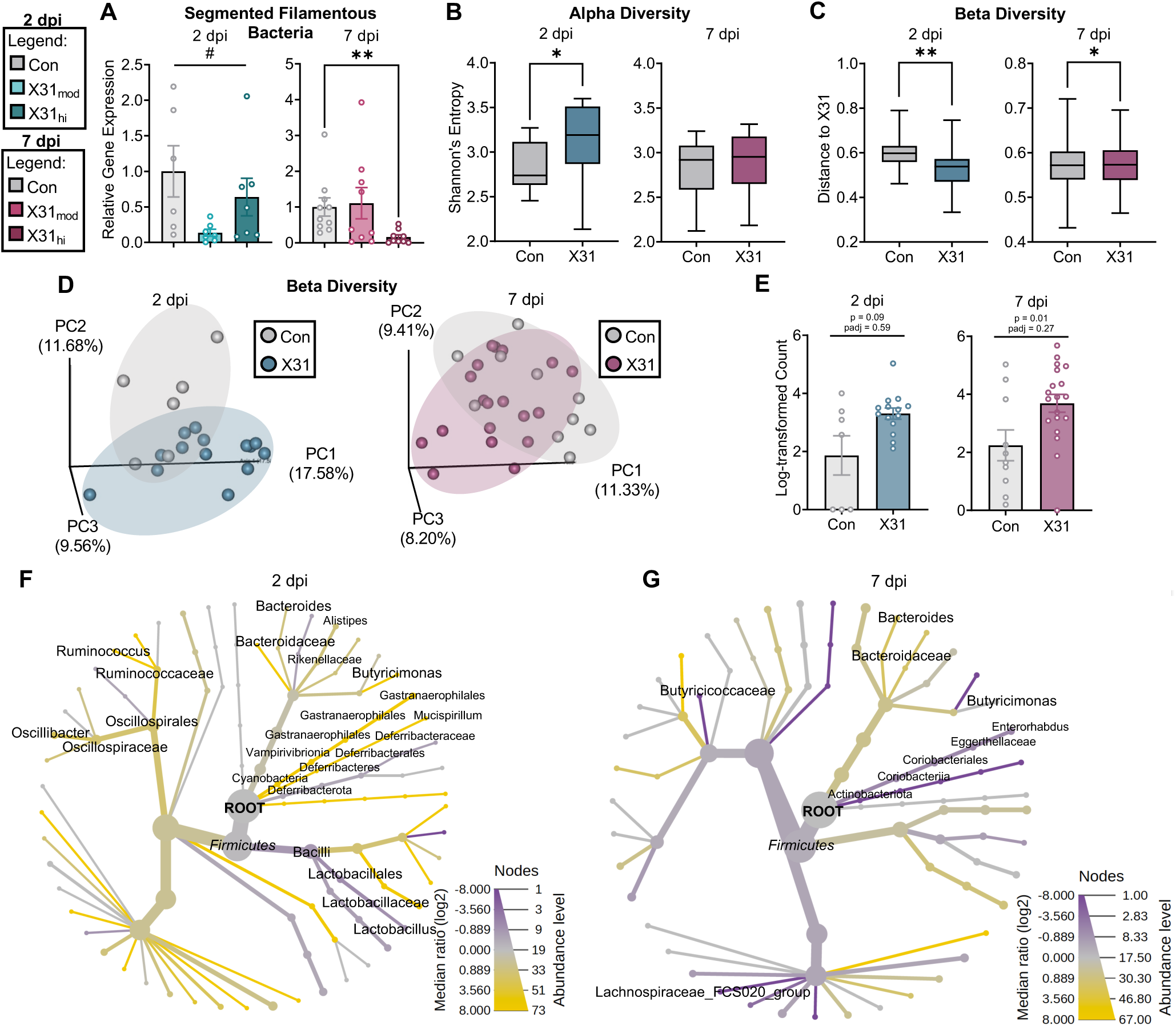
Respiratory IAV infection alters the maternal colonic microbiome. **(A)** Relative gene expression via qPCR showed a decrease in SFB, a bacterial regulator of T_H_17 cells, in the colon contents of high-dose dams at 7 dpi. **(B)** Alpha diversity (within-sample diversity) was different between con and X31 dams at 2 dpi only. **(C)** Beta diversity (between-sample diversity) was different between con and X31 dams at 2 and 7 dpi. **(D)** PCA based on unweighted UniFrac distance metrics. The ovals in the figure do not represent any statistical significance but serve as a visual guide to group differences. **(E)** Centered log-ratio analysis revealed a trending increase in *Bacteroides* in IAV-infected dams at both time points (DESeq2, padj < 0.05). **(F-G)** Heat tree analysis at **(F)** 2 and **(G)** 7 dpi revealed taxonomic differences between microbial communities (Wilcoxon Rank Sum test; italicized = not significant). *PCA* = principal component analysis, *IAV* = influenza A virus, *dpi* = days post-inoculation, *Con* = saline control, *X31_mod_* = IAV-X31 10^3^ TCID_50_, *X31_hi_* = IAV-X31 10^4^ TCID_50_, *X31* = X31_mod_ and X31_hi_. Shannon’ Index was used to calculate alpha diversity and unweighted UniFrac was used to calculate beta diversity. Data are means ± SEM; # = p < 0.1, * = p < 0.05, ** = p < 0.01; dots represent individual dams; n = 7-10 per treatment group.

Intestinal microbial dysbiosis as a characteristic of respiratory IAV infection has been found in both human and animal models [23,52–54]. To evaluate potential microbial disruption, we performed 16S rRNA sequencing on the colon contents of control and IAV-infected mice at 2 and 7 dpi. There was no difference between X31_mod_ and X31_hi_ treatment groups when evaluating within-sample diversity (alpha diversity) or between-sample diversity (beta diversity) at 2 dpi (p- and q-value = 0.75; p- and q-value =0.25, respectively) or 7 dpi (p- and q-value = 0.93; p- and q-value = 0.74, respectively); therefore, X31 treatment groups were collapsed. Alpha diversity was only different at 2 dpi (**Fig. 3B**) whereas beta-diversity was different at both endpoints (**Fig. 3C-D**). Taxonomic analysis revealed no differentially expressed bacteria at the genus level when adjusting for false discovery rate. However, there was a trending increase in *Bacteroides* across gestational IAV infection (**Fig. 3E**). Heat tree analysis verified the upregulation in *Bacteroides* along with the downregulation of several genus-level microbes associated with the Firmicutes phylum (**Fig. 3F-G**). A decrease in Firmicutes and upregulation in Bacteroidetes (the phyla *Bacteroides* falls under) was previously reported in mice with respiratory syncytial virus (RSV) infection [53] and in humans with IAV infection [55]. Altogether, these data verify colonic microbial alterations in gestating dams upon exposure to IAV.

### Gestational IAV infection leads to cortical abnormalities in the fetal brain

We next wanted to see if maternal IAV infection led to cortical abnormalities as previously described in mimetic-induced MIA models [20,21,26,56–59]. We performed immunohistochemistry on fetal brains at E16.5, as cortical layers are not distinguishable until E13.5 [34]. We examined the primary somatosensory cortex (**Fig. 4A**) based on prior MIA studies [58]. We observed a decrease in layer II/III upper excitatory neurons (SATB2 mean fluorescence intensity, MFI; **Fig. 4B**) in fetal brains from high-dose infected dams, with a trending decrease in layer IV-V deep excitatory neurons (TBR1 MFI; **Fig. 4C**), which is consistent with prior studies [21]. These group differences were apparent in both the right and left hemispheres (**Fig. S5A-B**) and did not differ between hemispheres (**Fig. S5C-D)**. We also saw a dose-dependent decrease in cortical plate thickness (**Fig. 4D**), a pathological indication of NDDs in humans [60–62]. We then divided this somatosensory cortical region of interest (ROI) into 10 equal laminar bins to determine the presence or absence of cortical malformations as previously described [21,57–59]. Significant differences were observed in layers 1, 7, and 8 for SATB2 MFI (**Fig. 4E**) and layers 3, 9, and 10 for TBR1 MFI in a dose-dependent manner (**Fig. 4F**). Overall, these data indicate that prenatal respiratory IAV infection leads to perturbations in fetal cortical development in a dose-dependent manner, providing further support for the existence of an infection severity threshold.

**Figure 4.**
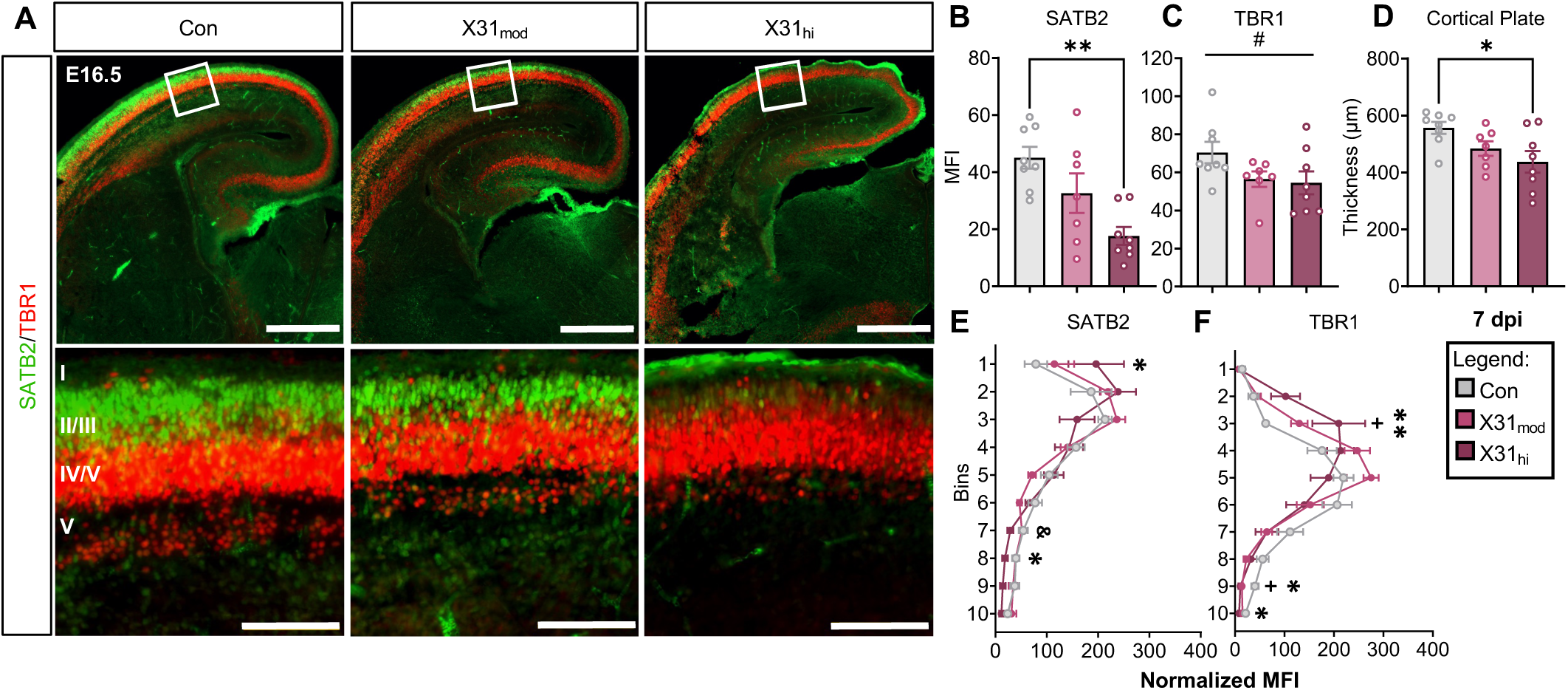
Respiratory IAV infection during pregnancy impacts cortical development in fetal brains from high-but not moderate-dose dams at 7 dpi. **(A)** Representative images of E16.5, 7 dpi fetal brains stained for SATB2 (green) and TBR1 (red). Close-up images were taken in the right-hemisphere somatosensory cortex in a 300 x 300 µm^2^ ROI. Top scale bars = 500µm; bottom scale bars = 100µm. I-V in the bottom left image represents cortical layers 1-6 **(B)** MFI of SATB2, an upper excitatory neuronal marker, is decreased in the fetal brains of high-dose mothers. **(C)** MFI of TBR1, a deep excitatory neuronal marker, tended to be decreased across all three groups. **(D)** Prenatal exposure to IAV-X31 reduced cortical thickness in fetal brains from X31_hi_ dams. **(E-F)** Dividing the ROI into 10 equal cortical laminar bins showed altered cortical lamination in **(E)** SATB2 bins 1, 7, and 8 and **(F)** TBR1 bins 3, 9, and 10 (* = Con vs X31_hi_, & = X31_mod_ vs X31_hi_, + = Con vs X31_mod_). *IAV* = influenza A virus, *dpi* = days post-inoculation, *E* = embryonic day, *Con* = saline control, *X31_mod_* = IAV-X31 10^3^ TCID_50_, *X31_hi_* = IAV-X31 10^4^ TCID_50,_ *ROI* = region of interest, *MFI* = mean fluorescence intensity. Groups were compared with one-way ANOVA with Tukey post hoc for multiple comparisons. For data containing residuals with unequal variance, Brown-Forsythe and Welch’s ANOVA with Dunnett T3 post hoc multiple comparisons was used. For non-parametric data, Kruskal-Wallis ANOVA with Dunn’s correction for multiple comparisons was used. Data are means ± SEM; # = p < 0.1, one symbol = p < 0.05, two symbols = p < 0.01; dots represent one representative fetus per litter; n = 9-10 per treatment group.

### Bulk RNA sequencing of the fetal brain reveals IAV-dependent transcriptional changes related to synaptic signaling and neuronal development

Since a high dose of IAV was required to produce cortical abnormalities, we performed bulk RNA-sequencing on control and X31_hi_ fetal brains at E16.5, 7 dpi. Differential gene expression analysis revealed 211 differentially upregulated and 173 differentially downregulated genes (**Fig. S1D**). We then used DAVID to determine the top ten significantly enriched up- and down-regulated GO pathways. Upregulated GO pathways largely corresponded to synaptic signaling (**Fig. 5A**), and downregulated GO pathways largely corresponded to neuronal and cellular development (**Fig. 5B; Supp. Table S8**). These top GO pathways are in alignment with GO pathways enriched in human NDDs [63,64]. Furthermore, a myriad of differentially expressed genes in the selected pathways are candidate genes for NDDs based on the Developmental Brain Disorder Gene Database (DBD) and the Simons Foundation Autism Research Initiative (SFARI) database (**Fig. 5C-D**, as indicated by * and + symbols, respectively). Notably, transcription of *Il17ra* and *Il17rc* (encoding the receptor complex for IL-17A and F) was unchanged in X31_hi_ fetal brains (adj. p = 0.61 and 0.83, respectively). Furthermore, no transcriptional differences were observed for *Il6ra* (adj. p = 0.84) or interferon-related genes (**Supp. Table S9**).

**Figure 5.**
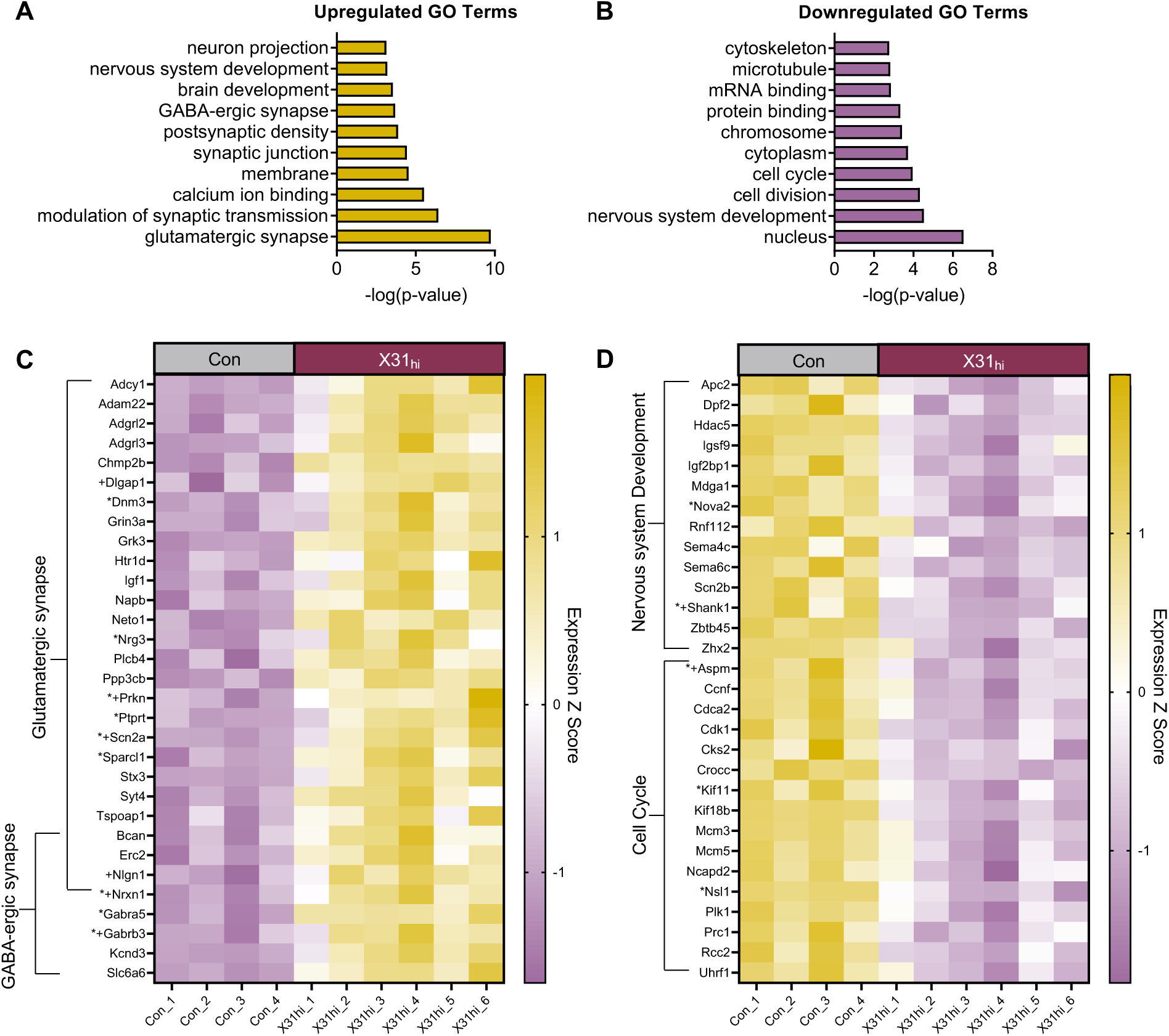
Bulk RNA-seq reveals genes enriched in neuronal development and synaptic signaling in fetal brains exposed to a high dose of prenatal IAV infection at E16.5, 7 dpi. Top ten significantly enriched **(A)** upregulated and **(B)** downregulated GO pathways in fetal brains prenatally exposed to a high dose of IAV, quantified by –log (p-value). Pathways were generated from the Database for Annotation, Visualization, and Integrated Discovery terms (DAVID). **(C)** Heatmap of genes differentially upregulated in the glutamatergic and GABA-ergic synapse pathways. **(D)** Heatmap of genes differentially downregulated in the nervous system development and cell cycle pathways. Genes with * represent candidate genes for neurodevelopmental disorders based on the Developmental Brain Disorder Gene Database (DBD). Genes with + represent candidate genes for Autism Spectrum Disorder based on the Simons Foundation Autism Research Initiative (SFARI) genes database. *IAV* = influenza A virus, *dpi* = days post inoculation, *Con* = saline control, *X31_hi_* = IAV-X31 10^4^ TCID_50,_ *GO* = gene ontology. Benjamini-Hochberg’s correction for false discovery (p < 0.1) was used to identify differentially expressed genes. Data are standardized logCPM values (Z score); dots represent one representative fetus per litter; n = 4-6 per treatment group.

The glutamatergic synapse pathway (GO:0098978) was the most significantly enriched upregulated GO term (**Fig. 5A**). Excitatory synaptic cell-adhesion molecules, *Nrxn1* and *Nlgn1,* were upregulated along with a trending upregulation in the gene encoding for excitatory presynaptic glutamate transporter, VGLUT2 (*Slc17a6;* adj. p = 0.17). Interestingly, genes encoding for excitatory postsynaptic scaffolding protein SHANK3 (*Shank3*; adj. p = 0.09) and glutamatergic postsynaptic density protein 95 (*Dlg4;* adj. p = 0.16), trended towards downregulation. This may indicate an imbalance in the development of pre- and post-synaptic terminals during gestational IAV infection. The GABA-ergic synapse pathway (GO:0098982) was also upregulated, as evidenced by upregulation in GABA receptor genes *Gabrb3 and Gabra5* (**Fig. 5C**).

Nervous system development (GO:0007399) and cell cycle (GO:0007049) were among the most significantly enriched downregulated GO pathways (**Fig. 5B**). This was evident in the downregulation of genes encoding for proteins related to cortical lamination, like *Apc2* and *Mdga1* (**Fig. 5D**). *Apc2* is involved in neuronal migration and axonal projection [65], and *Mdga1* plays a role in upper excitatory neuronal migration [66]. Genes responsible for proper embryonic neuronal migration and development including *Aspm* [67], *Cdk1* [68], and *Cks2* [69]—each of which are also involved in cell cycling—were also downregulated (**Fig. 5D**). Notably, transcription of *Satb2* and *Tbr1* was unchanged (adj. p = 0.61 and 0.49, respectively). Overall, these data indicate that gene pathways related to synaptic signaling and neuronal development are among the most dysregulated in the developing brain following prenatal exposure to a high dose of IAV.

### Gestational IAV infection alters embryonic border-associated macrophages—but not microglia— in a dose- and time-dependent manner

While some genetic mutations may underly cortical malformations [70,71], exactly what leads to perturbations in neocortical development during prenatal inflammation remains undetermined. Several studies implicate fetal microglia in MIA-related pathology [25,30,72–75]. Here, we hypothesized that an IAV immune insult would redirect microglia from their normal neurotrophic support functions, leading to neocortical abnormalities. To differentiate between microglia and BAMs, we co-stained fetal brain sections with CD206 and Iba1 at E11.5 and 16.5 (**Fig. 6A & F**). We observed no differences in number of microglia (CD206^-^Iba1^+^) across the whole brain at 2 (**Fig. 6B**) or 7 dpi (**Fig. 6D**). When assessed within and across specific fore-, mid-, and hind-brain regions (E11.5; **Fig. S6A**) or between the left and right hemispheres (E16.5; **Fig. S6B**), microglia density remained unchanged.

**Figure 6.**
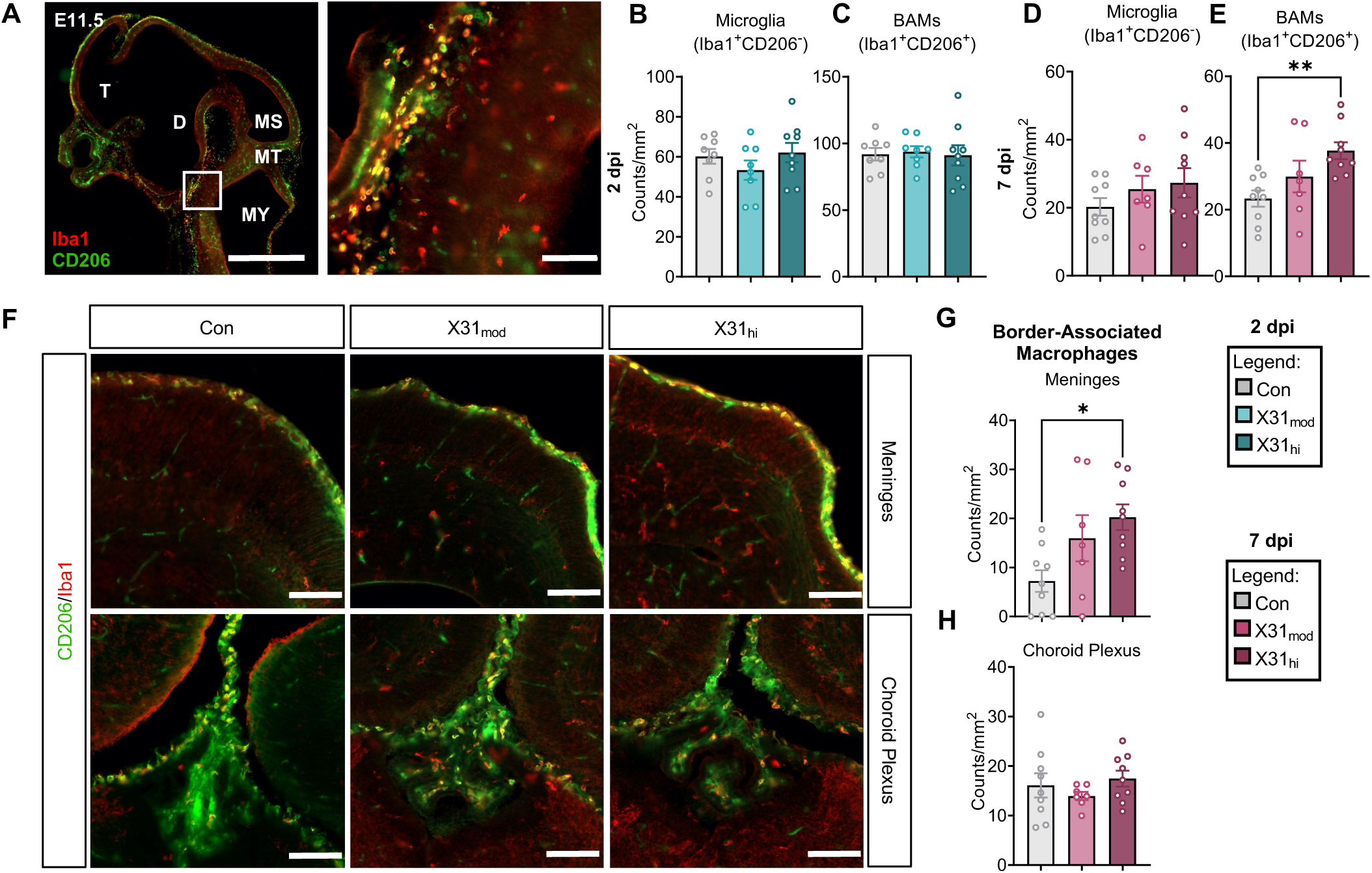
Border-associated macrophages but not microglia are upregulated in fetal brains from high-dose IAV dams. **(A)** Representative sagittal sections of the E11.5 fetal brain stained with Iba1 (red) and CD206 (green). Microglia are Iba1^+^CD206^-^ and BAMs are Iba1^+^CD206^+^. Left scale bar = 1000µm; right scale bar = 100µm. There were no changes in **(B)** microglia or **(C)** BAM count per mm^2^ at E11.5, 2 dpi. While there were no changes in **(D)** microglia count, **(E)** BAM count was upregulated in E16.5, 7 dpi fetal brains exposed to a high dose of IAV. **(F)** Representative coronal sections of the meninges and choroid plexus from each treatment group of Iba1 and CD206 stained E16.5 fetal brains. All scale bars = 100µm. **(G-H)** Separation of choroid plexus and meningeal BAMs showed that only **(H)** meningeal BAMs were increased at 7 dpi. *IAV* = influenza A virus, *dpi* = days post-inoculation, *E* = embryonic day, *Con* = saline control, *X31_mod_* = IAV-X31 10^3^ TCID_50_, *X31_hi_* = IAV-X31 10^4^ TCID_50_, *MFI* = mean fluorescence intensity, *T* = telencephalon, *D* = diencephalon, *MS* = mesencephalon, *MT* = metencephalon *MY* = myencephalon. Groups were compared using one-way ANOVA with Tukey post hoc for multiple comparisons. Data are means ± SEM; * = p < 0.05, ** = p < 0.01; dots represent one representative fetus per litter; n = 9-14 per treatment group.

Little is known about BAMs under homeostatic conditions, and even less is known about their function during embryonic development [76]. However, these transcriptionally distinct macrophages migrate from the yolk sac around the same time as microglia and take up residence at the brain’s borders [77,78]. Therefore, it is possible they also play a role in neuronal development. In line with this hypothesis, we observed an increase in BAMs (CD206^+^Iba1^+^) in fetal brains from X31_hi_ dams (**Fig. 6E**), which manifested at 7 dpi but was not apparent at 2 dpi (**Fig. 6C, S6C**). This observation was not restricted to a specific hemisphere (**Fig. S6D**). Previous studies have found upregulation in choroid plexus (ChP) BAMs in poly I:C-MIA fetal brains [26]. While we did not observe a difference in ChP BAMs (**Fig. 6H**), meningeal BAMs were significantly upregulated in fetal brains from high-dose IAV dams (**Fig. 6G**). Interestingly, *Mrc1*, the gene encoding for CD206, was not altered in the sequencing data set (adj. p = 0.88).

We then assessed whether or not prenatal IAV infection altered the proliferative capacity of Iba1^+^ cells, as seen in other models [25,79]. Cell proliferation, as measured by co-expression with nuclear Ki67 (**Fig. S6E**), was not altered at either endpoint (**Fig. S6F, H**) and did not change based on the brain region evaluated (**Fig. S6J-K**). Furthermore, the fluorescence intensity of Ki67-positive staining did not differ (**Fig. S6G, I, L-M**). Overall, these data demonstrate that despite a lack of changes in embryonic microglia, BAM numbers are elevated in the developing brain during late-stage IAV infection.

## Discussion

The United States has seen a significant increase in the incidence of NDDs over the past two decades [80]. As genetic risk factors comprise a small percentage of NDD etiologies, it is important to evaluate additional risk factors. In the 1990s, S.H. Fatemi and colleagues developed a mouse model of prenatal exposure to a neurotropic strain of IAV to evaluate the epidemiological link between IAV and offspring NDDs [11–13]. Since then, the majority of animal studies have used immunostimulants to identify potential mechanisms of gestational maternal inflammation. In this study, we use a non-neurotropic strain of IAV that recapitulates the severity of seasonal influenza infections in humans [81]. Critically, our work reveals that an infection severity threshold exists in tissues downstream of the site of infection. While IAV dose did not impact maternal lung lesions or viral transcripts, it played a significant role in fetal brain abnormalities. IAV infection severity also dictated downstream maternal intestinal immune dysfunction, which coincided with mild but significant T_H_17 cell suppression.

IAV infection can trigger adaptive immune responses that arise from a disruption in endogenous microbes, which appear to be at the crux of IAV-induced intestinal inflammation (as reviewed in [82]). Gastroenteritis-like symptoms, such as colonic shortening and shifts in the intestinal microbiome, are often evident during respiratory IAV infection even though the virus does not infect intestinal tissue [52]. Importantly, microbe depletion prior to IAV inoculation has been shown to diminish IAV-induced IL-17A production and intestinal injury [23]. Overall, the evidence suggests that microbial disruption precedes intestinal inflammation during respiratory IAV infection, leading to the differentiation of naïve CD4^+^ cells into pathogenic T_H_17 cells [48]. More importantly, infection-induced dysbiosis is a clinically relevant phenotype that is not fully recapitulated in poly I:C MIA models, wherein pre-existing homeostatic T_H_17 cells are indirectly (and more immediately) activated following poly I:C recognition by dendritic cells [20]. While our data indicate persistent colonic shortening throughout IAV replication and changes in microbial composition, we observe reduced production of IL-17 and lower numbers of intestinal T_H_17 cells post-IAV infection. Although this suggests a deviation from previously published observations with highly pathogenic IAV strain PR8 [23], it is possible that IL-17 production could be elevated at some point between 2 and 7 dpi in our X31 model. It is also possible that altered immune states during pregnancy lead to a unique intestinal immune phenotype following respiratory IAV infection, although this remains to be tested.

Furthermore, while our flow cytometry data do not indicate a pathogenic T_H_17 phenotype, our gene expression data indicate elevated IL-6 and decreased TGF-β in the maternal intestine. T_H_17 cells can be primed towards a pathogenic phenotype in the absence of *Tgfb1* and the presence of *Il6* and *Il23a* [49,83]. Thus, intestinal T_H_17-cell priming antigen-presenting cells will also need to be evaluated to determine whether they can generate pathogenic intestinal T_H_17 cells at alternate time points in this model.

In poly I:C-induced MIA models, pharmacological inhibition and genetic knockouts of maternal IL-17A are sufficient to halt the development of offspring cortical patches and behavioral deficits [21]. These offspring abnormalities are also absent in poly I:C-challenged animals that do not harbor SFB, and thus do not have sufficient populations of T_H_17 cells to mount an IL-17 response [20,22]. Contrary to these findings, we observe cortical abnormalities in our model of IAV-induced MIA despite persistent downregulation of intestinal IL-17 responses and SFB transcripts. Notably, while anti-viral IL-17 responses in maternal lungs were within normal ranges for IAV infection, circulating IL-17 levels remained low and did not differ between groups. This suggests that the alterations in neocortical development observed here are inducible via IL-17-independent mechanisms. While further testing is required to parse this out, it is important to note the consistent elevation in IL-6 across maternal tissue types (lung, serum, intestine) in our model. Indeed, the idea that maternal IL-6 mediates offspring neurodevelopmental abnormalities predates the more recent interest in IL-17A. Early poly I:C-induced MIA studies revealed that pharmacological inhibition and genetic depletion of IL-6 prevented certain aspects of behavioral deficits in offspring [84]. Genetic ablation of the IL-6 receptor on placental trophoblasts in poly I:C-challenged dams prevented offspring behavioral abnormalities and neuropathologies [85]. Recent studies further bolster the idea that IL-6 is capable of directly mediating brain development via prenatal programming of synaptogenesis (as seen in mice) and via dysregulation of radial glia (as seen in human brain organoids) [86,87]. There is also strong evidence documenting the negative effects of anti-viral interferon signaling on placental and fetal development during maternal infection [88]. The type I and II interferon signaling patterns observed in our model may be acting via similar mechanisms. Therefore, it is possible that maternal IL-6 and/or IFN signaling—rather than IL-17—could be driving downstream intestinal inflammation and disrupted neocortical phenotypes in our model.

Alterations in neocortical development can have life-long negative consequences for behavior and neural processing, and disordered cortical lamination is common in NDDs [89]. Here, we observed a reduction in superficial SATB2^+^ neurons in the neocortex of E16.5 fetuses exposed to high but not moderate dose maternal IAV infection. Several mimetic-induced MIA models report the same reduction in fetal SATB2^+^ neurons and improper cortical layering of SATB2 and TBR1 before the onset of behavioral deficits [21,57,59,90], suggesting that this phenomenon is conserved across disparate models of MIA. The importance of SATB2 can be appreciated when observing patients with genetic mutations and deletions of *Satb2*. This SATB2-associated syndrome causes a myriad of complications, including autistic-like behavior and speech impairments [91]. Interestingly, influenza vaccination in early pregnancy has been shown to prevent a reduction in embryonic SATB2 in a rodent model of LPS-induced MIA [59]. While it stands to reason that maternal vaccination against IAV before infection would similarly protect against SATB2 abnormalities in our model, this still needs to be tested. Although others have proposed that upregulation of IL-17 receptors on embryonic neurons precedes SATB2 abnormalities [58], we observed no differences in IL-17 receptor expression in the embryonic brain. Overall, SATB2^+^ neurons appear to be conserved cellular targets of maternal inflammation and are likely disrupted through an IL-17-independent mechanism in our model.

Our RNA-seq data from X31_hi_ fetal brains bolsters the evidence for altered cortical lamination. For instance, we see a reduction in genes *Apc2, Mdga1,* and *Usp11.* Genetic knockout or loss of function studies performed on each of these genes reveal their importance in regulating early neuronal layering in the murine cortex. Knockout of *Apc2* leads to improper cortical lamination of TBR1^+^ neurons at E16.5 [65]. Knockout of *Usp11*, which encodes a protein responsible for cortical neurogenesis and migration, leads to reduced migration of neuronal progenitors into superficial layers (e.g., reduction in SATB2 in layers II-IV) [92]. Loss of function of *Mdga1* indicated that this gene is critical for radial migration of upper layer neurons [66]. Overall, the downregulation of these genes, among others, supports the idea that severe maternal IAV infection disrupts processes related to neocortical formation and function.

As cortical neurogenesis precedes synaptogenesis [28], alterations in cortical lamination are often accompanied by dysregulated synaptic signaling [93,94]. The observed upregulation in glutamatergic synapse-related genes in our model corroborates evidence from a recent study demonstrating that direct intracerebroventricular injection of IL-6 into the embryonic brain elevates genetic programs of synaptogenesis in cortical glutamatergic neurons [86]. Poly I:C-induced MIA studies also describe upregulated glutamatergic synapse density in cortical upper layer [95] and deep layer [96] neurons. Critically, our data are also in alignment with clinical studies that demonstrate dysregulation in both cortical lamination and synapse development and function (e.g., excitatory/inhibitory imbalance) in patients with NDDs [89,97]. Future studies are needed to determine whether IAV-induced changes in fetal brain organization and developmental processing persist postnatally and whether they correspond with NDD-related behaviors.

We hypothesized that microglia and BAMs would be prime cellular candidates for orchestrating cortical pathologies. At the time of our maternal IAV challenge, microglia are migrating to the embryonic brain, where they aid in critical neurotrophic support functions including cortical development [29,30] and cortical synaptic formation [98]. The role microglia play in NDDs is debated; however, numerous studies report that microglia are impacted by MIA [27,74,99]. While our histological analysis of embryonic microglia indicates that their migration patterns and proliferation are not altered by maternal IAV infection, functional assays are needed to confirm whether microglial activities are shifted in the immediate days following infection. Indeed, microglial density and proliferative capacity may not directly correlate with phagocytic function nor with the production of neurotrophic or inflammatory mediators, each of which has been shown to modulate critical neurodevelopmental processes (as reviewed in [19]).

Interestingly, the dose-dependent increased density of BAMs indicates a macrophage-specific response to IAV-induced maternal inflammation. While BAMs do not infiltrate the brain parenchyma to interact with early neuronal cells like microglia do, they are in direct contact with the periphery and thus play a critical role in immune defense [77,100]. At least one study demonstrates that poly I:C-induced MIA directly impacts the homeostatic function of embryonic brain macrophages, leading to increases in ChP macrophage populations [26]. These authors propose that an accumulation of macrophages at the ChP and ventricular zone indirectly contributes to the cortical malformations observed in their model, whereby increased recruitment of phagocytes into the brain parenchyma disrupts neural progenitor proliferation at the cortex [26]. Direct examination of this proposed mechanism, however, is still needed. Notably, we observed an increased abundance of CD206^+^ macrophages in the meninges but not the ChP in our model. To our knowledge, no study has specifically evaluated embryonic meningeal BAMs following maternal inflammation, nor has any study compared BAM subsets in this context. Further work is needed to elucidate the role BAMs might play in regulating prenatal neuronal patterning following maternal IAV infection and the mechanisms by which they might fulfill these roles.

Taken together, our data suggest that IAV-induced cortical malformations and altered macrophage populations in the embryonic brain arise independently of maternal IL-17 signaling. Importantly, we demonstrate that a higher infectious dose of IAV is necessary to induce downstream changes in the maternal intestine and the fetal brain, further bolstering our previous findings [31]. Overall, the discrepancies and similarities we observed between our model of live IAV-induced MIA and models of mimetic-induced MIA highlight the importance of using live pathogens to evaluate the complete immune response and to improve translation to the clinic.

## Supporting information

MIA Reporting Guidelines

## Acknowledgments

We would like to thank Dr. Jacob Yount for providing the IAV viral strain; Dr. Shih-Hsuan Hsiao for scoring lung lesions; Dr. Alvaro Hernandez, Dr. Jenny Drnevich, and Dr. Chris Wright for RNA-Sequencing processing; and Izan Chalen for helping with tissue collection. This study was supported by the Roy J. Carver Charitable Trust (Grant #23-5683) and by start-up funds from the University of Illinois Urbana-Champaign to A.M.A. Figure 1 schematic was generated with Biorender.com.

## CRediT Author Contributions

**Ashley M. Otero**: Conceptualization, Methodology, Validation, Formal Analysis, Investigation, Writing – Original Draft, Writing – Review & Editing, Visualization, Project Administration. **Meghan G. Connolly**: Methodology, Formal Analysis, Investigation, Writing – Review & Editing. **Rafael J. Gonzalez-Ricon**: Investigation. **Selena S. Wang**: Investigation. **Jacob M. Allen**: Conceptualization, Software, Writing – Review & Editing. **Adrienne M. Antonson**: Conceptualization, Methodology, Writing – Review & Editing, Supervision, Funding Acquisition.

## References

1. Oseghale O, Vlahos R, O’Leary JJ, Brooks RD, Brooks DA, Liong S, et al. Influenza Virus Infection during Pregnancy as a Trigger of Acute and Chronic Complications. Viruses. 2022 Dec 7;14(12):2729.

2. Raj RS, Bonney EA, Phillippe M. Influenza, Immune System, and Pregnancy. Reprod Sci. 2014 Dec;21(12):1434–51.

3. Mertz D, Geraci J, Winkup J, Gessner BD, Ortiz JR, Loeb M. Pregnancy as a risk factor for severe outcomes from influenza virus infection: A systematic review and meta-analysis of observational studies. Vaccine. 2017 Jan 23;35(4):521–8.

4. Dawood FS, Kittikraisak W, Patel A, Rentz Hunt D, Suntarattiwong P, Wesley MG, et al. Incidence of influenza during pregnancy and association with pregnancy and perinatal outcomes in three middle-income countries: a multisite prospective longitudinal cohort study. The Lancet Infectious Diseases. 2021 Jan;21(1):97–106.

5. Abu-Raya B, Michalski C, Sadarangani M, Lavoie PM. Maternal Immunological Adaptation During Normal Pregnancy. Front Immunol. 2020 Oct 7;11:575197.

6. Kępińska AP, Iyegbe CO, Vernon AC, Yolken R, Murray RM, Pollak TA. Schizophrenia and Influenza at the Centenary of the 1918-1919 Spanish Influenza Pandemic: Mechanisms of Psychosis Risk. Front Psychiatry. 2020 Feb 26;11:72.

7. Brown AS, Begg MD, Gravenstein S, Schaefer CA, Wyatt RJ, Bresnahan M, et al. Serologic Evidence of Prenatal Influenza in the Etiology of Schizophrenia. Arch Gen Psychiatry. 2004 Aug 1;61(8):774.

8. Brown AS, Patterson PH. Maternal Infection and Schizophrenia: Implications for Prevention. Schizophrenia Bulletin. 2011 Mar 1;37(2):284–90.

9. Parboosing R, Bao Y, Shen L, Schaefer CA, Brown AS. Gestational Influenza and Bipolar Disorder in Adult Offspring. JAMA Psychiatry. 2013 Jul 1;70(7):677.

10. Atladóttir HÓ, Thorsen P, Østergaard L, Schendel DE, Lemcke S, Abdallah M, et al. Maternal Infection Requiring Hospitalization During Pregnancy and Autism Spectrum Disorders. J Autism Dev Disord. 2010 Dec;40(12):1423–30.

11. Fatemi SH, Emamian ES, Sidwell RW, Kist DA, Stary JM, Earle JA, et al. Human influenza viral infection in utero alters glial fibrillary acidic protein immunoreactivity in the developing brains of neonatal mice. Mol Psychiatry. 2002 Jul 1;7(6):633–40.

12. Fatemi SH, Pearce DA, Brooks AI, Sidwell RW. Prenatal viral infection in mouse causes differential expression of genes in brains of mouse progeny: A potential animal model for schizophrenia and autism. Synapse. 2005 Aug;57(2):91–9.

13. Fatemi SH, Reutiman TJ, Folsom TD, Huang H, Oishi K, Mori S, et al. Maternal infection leads to abnormal gene regulation and brain atrophy in mouse offspring: Implications for genesis of neurodevelopmental disorders. Schizophrenia Research. 2008 Feb;99(1–3):56–70.

14. Shi L, Tu N, Patterson PH. Maternal influenza infection is likely to alter fetal brain development indirectly: the virus is not detected in the fetus. Int j dev neurosci. 2005 Apr;23(2–3):299–305.

15. Meyer U. Prenatal Poly(I:C) Exposure and Other Developmental Immune Activation Models in Rodent Systems. Biological Psychiatry. 2014 Feb;75(4):307–15.

16. Meyer U, Feldon J, Fatemi SH. In-vivo rodent models for the experimental investigation of prenatal immune activation effects in neurodevelopmental brain disorders. Neuroscience & Biobehavioral Reviews. 2009 Jul;33(7):1061–79.

17. Braciale TJ, Sun J, Kim TS. Regulating the adaptive immune response to respiratory virus infection. Nat Rev Immunol. 2012 Apr;12(4):295–305.

18. Iwasaki A, Pillai PS. Innate immunity to influenza virus infection. Nat Rev Immunol. 2014 May;14(5):315–28.

19. Otero AM, Antonson AM. At the crux of maternal immune activation: Viruses, microglia, microbes, and IL-17A. Immunological Reviews. 2022 Oct;311(1):205–23.

20. Kim S, Kim H, Yim YS, Ha S, Atarashi K, Tan TG, et al. Maternal gut bacteria promote neurodevelopmental abnormalities in mouse offspring. Nature. 2017 Sep 28;549(7673):528–32.

21. Choi GB, Yim YS, Wong H, Kim S, Kim H, Kim SV, et al. The maternal interleukin-17a pathway in mice promotes autism-like phenotypes in offspring. Science. 2016 Feb 26;351(6276):933–9.

22. Lammert CR, Frost EL, Bolte AC, Paysour MJ, Shaw ME, Bellinger CE, et al. Cutting Edge: Critical Roles for Microbiota-Mediated Regulation of the Immune System in a Prenatal Immune Activation Model of Autism. JI. 2018 Aug 1;201(3):845–50.

23. Wang J, Li F, Wei H, Lian ZX, Sun R, Tian Z. Respiratory influenza virus infection induces intestinal immune injury via microbiota-mediated Th17 cell–dependent inflammation. Journal of Experimental Medicine. 2014 Nov 17;211(12):2397–410.

24. Stockinger B, Omenetti S. The dichotomous nature of T helper 17 cells. Nat Rev Immunol. 2017 Sep;17(9):535–44.

25. Ben-Yehuda H, Matcovitch-Natan O, Kertser A, Spinrad A, Prinz M, Amit I, et al. Maternal Type-I interferon signaling adversely affects the microglia and the behavior of the offspring accompanied by increased sensitivity to stress. Mol Psychiatry. 2020 May;25(5):1050–67.

26. Cui J, Shipley FB, Shannon ML, Alturkistani O, Dani N, Webb MD, et al. Inflammation of the Embryonic Choroid Plexus Barrier following Maternal Immune Activation. Developmental Cell. 2020 Dec;55(5):617–628.e6.

27. Yu D, Li T, Delpech JC, Zhu B, Kishore P, Koshi T, et al. Microglial GPR56 is the molecular target of maternal immune activation-induced parvalbumin-positive interneuron deficits. Sci Adv. 2022 May 6;8(18):eabm2545.

28. Reemst K, Noctor SC, Lucassen PJ, Hol EM. The Indispensable Roles of Microglia and Astrocytes during Brain Development. Front Hum Neurosci. 2016 Nov 8;10.3389/fnhum.2016.00566.

29. Antony JM, Paquin A, Nutt SL, Kaplan DR, Miller FD. Endogenous microglia regulate development of embryonic cortical precursor cells. J Neurosci Res. 2011 Mar;89(3):286–98.

30. Cunningham CL, Martinez-Cerdeno V, Noctor SC. Microglia Regulate the Number of Neural Precursor Cells in the Developing Cerebral Cortex. Journal of Neuroscience. 2013 Mar 6;33(10):4216–33.

31. Antonson AM, Kenney AD, Chen HJ, Corps KN, Yount JS, Gur TL. Moderately pathogenic maternal influenza A virus infection disrupts placental integrity but spares the fetal brain. Brain, Behavior, and Immunity. 2021 Aug;96:28–39.

32. Patten AR, Fontaine CJ, Christie BR. A Comparison of the Different Animal Models of Fetal Alcohol Spectrum Disorders and Their Use in Studying Complex Behaviors. Front Pediatr. 2014 Sep 3;2:93.

33. Schouten M, van der Sluijs KF, Gerlitz B, Grinnell BW, Roelofs JJ, Levi MM, et al. Activated protein C ameliorates coagulopathy but does not influence outcome in lethal H1N1 influenza: a controlled laboratory study. Crit Care. 2010;14(2):R65.

34. Chen VS, Morrison JP, Southwell MF, Foley JF, Bolon B, Elmore SA. Histology Atlas of the Developing Prenatal and Postnatal Mouse Central Nervous System, with Emphasis on Prenatal Days E7.5 to E18.5. Toxicol Pathol. 2017 Aug;45(6):705–44.

35. Kaufman MH, Baldock R, Bard JBL, Davidson D, Morriss-Kay G, editors. Kaufman’s atlas of mouse development: with coronal sections. Supplement. Amsterdam: Academic Press; 2016. 331 p.

36. Schambra U. Prenatal Mouse Brain Atlas. Boston, MA: Springer US; 2008.

37. Kim E, Tran M, Sun Y, Huh JR. Isolation and analyses of lamina propria lymphocytes from mouse intestines. STAR Protocols. 2022 Jun;3(2):101366.

38. Allen JM, Mackos AR, Jaggers RM, Brewster PC, Webb M, Lin CH, et al. Psychological stress disrupts intestinal epithelial cell function and mucosal integrity through microbe and host-directed processes. Gut Microbes. 2022 Dec 31;14(1):2035661.

39. Peixoto L, Risso D, Poplawski SG, Wimmer ME, Speed TP, Wood MA, et al. How data analysis affects power, reproducibility and biological insight of RNA-seq studies in complex datasets. Nucleic Acids Res. 2015 Sep 18;43(16):7664–74.

40. Chen Y, Lun ATL, Smyth GK. From reads to genes to pathways: differential expression analysis of RNA-Seq experiments using Rsubread and the edgeR quasi-likelihood pipeline. F1000Res. 2016 Aug 2;5:1438.

41. Sego TJ, Aponte-Serrano JO, Ferrari Gianlupi J, Heaps SR, Breithaupt K, Brusch L, et al. A modular framework for multiscale, multicellular, spatiotemporal modeling of acute primary viral infection and immune response in epithelial tissues and its application to drug therapy timing and effectiveness. Peirce SM, editor. PLoS Comput Biol. 2020 Dec 21;16(12):e1008451.

42. Miao H, Hollenbaugh JA, Zand MS, Holden-Wiltse J, Mosmann TR, Perelson AS, et al. Quantifying the early immune response and adaptive immune response kinetics in mice infected with influenza A virus. J Virol. 2010 Jul;84(13):6687–98.

43. Major J, Crotta S, Llorian M, McCabe TM, Gad HH, Priestnall SL, et al. Type I and III interferons disrupt lung epithelial repair during recovery from viral infection. Science. 2020 Aug 7;369(6504):712–7.

44. Sahu U, Biswas D, Prajapati VK, Singh AK, Samant M, Khare P. Interleukin-17—A multifaceted cytokine in viral infections. Journal Cellular Physiology. 2021 Dec;236(12):8000–19.

45. Crowe CR, Chen K, Pociask DA, Alcorn JF, Krivich C, Enelow RI, et al. Critical Role of IL-17RA in Immunopathology of Influenza Infection. The Journal of Immunology. 2009 Oct 15;183(8):5301–10.

46. Zhou L, Ivanov II, Spolski R, Min R, Shenderov K, Egawa T, et al. IL-6 programs TH-17 cell differentiation by promoting sequential engagement of the IL-21 and IL-23 pathways. Nat Immunol. 2007 Sep;8(9):967–74.

47. Chassaing B, Aitken JD, Malleshappa M, Vijay-Kumar M. Dextran Sulfate Sodium (DSS)-Induced Colitis in Mice. CP in Immunology. 2014 Feb;104(1: 10.1002/0471142735.im1525s104).

48. Omenetti S, Bussi C, Metidji A, Iseppon A, Lee S, Tolaini M, et al. The Intestine Harbors Functionally Distinct Homeostatic Tissue-Resident and Inflammatory Th17 Cells. Immunity. 2019 Jul;51(1):77–89.e6.

49. Ghoreschi K, Laurence A, Yang XP, Tato CM, McGeachy MJ, Konkel JE, et al. Generation of pathogenic TH17 cells in the absence of TGF-β signalling. Nature. 2010 Oct;467(7318):967–71.

50. Ivanov II, Frutos R de L, Manel N, Yoshinaga K, Rifkin DB, Sartor RB, et al. Specific Microbiota Direct the Differentiation of IL-17-Producing T-Helper Cells in the Mucosa of the Small Intestine. Cell Host & Microbe. 2008 Oct;4(4):337–49.

51. Ivanov II, Atarashi K, Manel N, Brodie EL, Shima T, Karaoz U, et al. Induction of Intestinal Th17 Cells by Segmented Filamentous Bacteria. Cell. 2009 Oct;139(3):485–98.

52. Deriu E, Boxx GM, He X, Pan C, Benavidez SD, Cen L, et al. Influenza Virus Affects Intestinal Microbiota and Secondary Salmonella Infection in the Gut through Type I Interferons. Tsolis RM, editor. PLoS Pathog. 2016 May 5;12(5):e1005572.

53. Groves HT, Cuthbertson L, James P, Moffatt MF, Cox MJ, Tregoning JS. Respiratory Disease following Viral Lung Infection Alters the Murine Gut Microbiota. Front Immunol. 2018 Feb 12;9:182.

54. Minodier L, Charrel RN, Ceccaldi PE, Van Der Werf S, Blanchon T, Hanslik T, et al. Prevalence of gastrointestinal symptoms in patients with influenza, clinical significance, and pathophysiology of human influenza viruses in faecal samples: what do we know? Virol J. 2015 Dec;12(1):215.

55. Al Khatib HA, Mathew S, Smatti MK, Eltai NO, Pathan SA, Al Thani AA, et al. Profiling of Intestinal Microbiota in Patients Infected with Respiratory Influenza A and B Viruses. Pathogens. 2021 Jun 17;10(6):761.

56. Canales CP, Estes ML, Cichewicz K, Angara K, Aboubechara JP, Cameron S, et al. Sequential perturbations to mouse corticogenesis following in utero maternal immune activation. eLife. 2021 Mar 5;10:e60100.

57. Ben-Reuven L, Reiner O. Dynamics of cortical progenitors and production of subcerebral neurons are altered in embryos of a maternal inflammation model for autism. Mol Psychiatry. 2021 May;26(5):1535–50.

58. Shin Yim Y, Park A, Berrios J, Lafourcade M, Pascual LM, Soares N, et al. Reversing behavioural abnormalities in mice exposed to maternal inflammation. Nature. 2017 Sep 28;549(7673):482–7.

59. Wu Y, Qi F, Song D, He Z, Zuo Z, Yang Y, et al. Prenatal influenza vaccination rescues impairments of social behavior and lamination in a mouse model of autism. J Neuroinflammation. 2018 Dec;15(1):228.

60. Goldman AL, Pezawas L, Mattay VS, Fischl B, Verchinski BA, Chen Q, et al. Widespread Reductions of Cortical Thickness in Schizophrenia and Spectrum Disorders and Evidence of Heritability. Arch Gen Psychiatry. 2009 May 1;66(5):467.

61. Hardan AY, Libove RA, Keshavan MS, Melhem NM, Minshew NJ. A Preliminary Longitudinal Magnetic Resonance Imaging Study of Brain Volume and Cortical Thickness in Autism. Biological Psychiatry. 2009 Aug;66(4):320–6.

62. Chiappelli J, Kochunov P, Savransky A, Fisseha F, Wisner K, Du X, et al. Allostatic load and reduced cortical thickness in schizophrenia. Psychoneuroendocrinology. 2017 Mar;77:105–11.

63. Parenti I, Rabaneda LG, Schoen H, Novarino G. Neurodevelopmental Disorders: From Genetics to Functional Pathways. Trends in Neurosciences. 2020 Aug;43(8):608–21.

64. Rahnama M, Tehrani HA, Mirzaie M, Vahid Ziaee. Identification of key genes and convergent pathways disrupted in autism spectrum disorder via comprehensive bioinformatic analysis. Informatics in Medicine Unlocked. 2021;24:100589.

65. Shintani T, Takeuchi Y, Fujikawa A, Noda M. Directional Neuronal Migration Is Impaired in Mice Lacking Adenomatous Polyposis Coli 2. J Neurosci. 2012 May 9;32(19):6468–84.

66. Takeuchi A, O’Leary DDM. Radial Migration of Superficial Layer Cortical Neurons Controlled by Novel Ig Cell Adhesion Molecule MDGA1. J Neurosci. 2006 Apr 26;26(17):4460–4.

67. Buchman JJ, Durak O, Tsai LH. ASPM regulates Wnt signaling pathway activity in the developing brain. Genes Dev. 2011 Sep 15;25(18):1909–14.

68. Baffet AD, Hu DJ, Vallee RB. Cdk1 Activates Pre-mitotic Nuclear Envelope Dynein Recruitment and Apical Nuclear Migration in Neural Stem Cells. Developmental Cell. 2015 Jun;33(6):703–16.

69. Frontini M, Kukalev A, Leo E, Ng YM, Cervantes M, Cheng CW, et al. The CDK Subunit CKS2 Counteracts CKS1 to Control Cyclin A/CDK2 Activity in Maintaining Replicative Fidelity and Neurodevelopment. Developmental Cell. 2012 Aug;23(2):356–70.

70. Chung C, Yang X, Bae T, Vong KI, Mittal S, Donkels C, et al. Comprehensive multi-omic profiling of somatic mutations in malformations of cortical development. Nat Genet. 2023 Feb;55(2):209–20.

71. Barkovich AJ, Guerrini R, Kuzniecky RI, Jackson GD, Dobyns WB. A developmental and genetic classification for malformations of cortical development: update 2012. Brain. 2012 May;135(5):1348–69.

72. Rosin JM, Sinha S, Biernaskie J, Kurrasch DM. A subpopulation of embryonic microglia respond to maternal stress and influence nearby neural progenitors. Developmental Cell. 2021 May;56(9):1326–1345.e6.

73. Ozaki K, Kato D, Ikegami A, Hashimoto A, Sugio S, Guo Z, et al. Maternal immune activation induces sustained changes in fetal microglia motility. Sci Rep. 2020 Dec;10(1):21378.

74. Bolton JL, Short AK, Othy S, Kooiker CL, Shao M, Gunn BG, et al. Early stress-induced impaired microglial pruning of excitatory synapses on immature CRH-expressing neurons provokes aberrant adult stress responses. Cell Reports. 2022 Mar;38(13):110600.

75. Sasaki T, Tome S, Takei Y. Intraventricular IL-17A administration activates microglia and alters their localization in the mouse embryo cerebral cortex. Mol Brain. 2020 Dec;13(1):93.

76. Gerganova G, Riddell A, Miller AA. CNS border-associated macrophages in the homeostatic and ischaemic brain. Pharmacology & Therapeutics. 2022 Dec;240:108220.

77. Utz SG, See P, Mildenberger W, Thion MS, Silvin A, Lutz M, et al. Early Fate Defines Microglia and Non-parenchymal Brain Macrophage Development. Cell. 2020 Apr;181(3):557–573.e18.

78. Goldmann T, Wieghofer P, Jordão MJC, Prutek F, Hagemeyer N, Frenzel K, et al. Origin, fate and dynamics of macrophages at central nervous system interfaces. Nat Immunol. 2016 Jul;17(7):797–805.

79. Loayza M, Lin S, Carter K, Ojeda N, Fan LW, Ramarao S, et al. Maternal immune activation alters fetal and neonatal microglia phenotype and disrupts neurogenesis in mice. Pediatr Res. 2023 Apr;93(5):1216–25.

80. Maenner MJ, Shaw KA, Bakian AV, Bilder DA, Durkin MS, Esler A, et al. Prevalence and Characteristics of Autism Spectrum Disorder Among Children Aged 8 Years — Autism and Developmental Disabilities Monitoring Network, 11 Sites, United States, 2018. MMWR Surveill Summ. 2021 Dec 3;70(11):1–16.

81. Hayward AC, Fragaszy EB, Bermingham A, Wang L, Copas A, Edmunds WJ, et al. Comparative community burden and severity of seasonal and pandemic influenza: results of the Flu Watch cohort study. The Lancet Respiratory Medicine. 2014 Jun;2(6):445–54.

82. Chen CJ, Wu GH, Kuo RL, Shih SR. Role of the intestinal microbiota in the immunomodulation of influenza virus infection. Microbes and Infection. 2017 Dec;19(12):570–9.

83. Lee Y, Awasthi A, Yosef N, Quintana FJ, Xiao S, Peters A, et al. Induction and molecular signature of pathogenic TH17 cells. Nat Immunol. 2012 Oct;13(10):991–9.

84. Smith SEP, Li J, Garbett K, Mirnics K, Patterson PH. Maternal Immune Activation Alters Fetal Brain Development through Interleukin-6. Journal of Neuroscience. 2007 Oct 3;27(40):10695–702.

85. Wu WL, Hsiao EY, Yan Z, Mazmanian SK, Patterson PH. The placental interleukin-6 signaling controls fetal brain development and behavior. Brain, Behavior, and Immunity. 2017 May;62:11–23.

86. Mirabella F, Desiato G, Mancinelli S, Fossati G, Rasile M, Morini R, et al. Prenatal interleukin 6 elevation increases glutamatergic synapse density and disrupts hippocampal connectivity in offspring. Immunity. 2021 Nov;54(11):2611–2631.e8.

87. Sarieva K, Kagermeier T, Khakipoor S, Atay E, Yentür Z, Becker K, et al. Human brain organoid model of maternal immune activation identifies radial glia cells as selectively vulnerable. Mol Psychiatry. 2023 Mar 6;10.1038/s41380-023-01997–1.

88. Yockey LJ, Iwasaki A. Interferons and Proinflammatory Cytokines in Pregnancy and Fetal Development. Immunity. 2018 Sep;49(3):397–412.

89. Guarnieri FC, De Chevigny A, Falace A, Cardoso C. Disorders of neurogenesis and cortical development. Dialogues in Clinical Neuroscience. 2018 Dec 31;20(4):255–66.

90. Carpentier PA, Haditsch U, Braun AE, Cantu AV, Moon HM, Price RO, et al. Stereotypical Alterations in Cortical Patterning Are Associated with Maternal Illness-Induced Placental Dysfunction. J Neurosci. 2013 Oct 23;33(43):16874–88.

91. Zarate YA, Kaylor J, Fish J. SATB2-Associated Syndrome. In: Adam MP, Feldman J, Mirzaa GM, Pagon RA, Wallace SE, Bean LJ, et al., editors. GeneReviews®. Seattle (WA): University of Washington, Seattle; 1993.

92. Chiang SY, Wu HC, Lin SY, Chen HY, Wang CF, Yeh NH, et al. Usp11 controls cortical neurogenesis and neuronal migration through Sox11 stabilization. Sci Adv. 2021 Feb 12;7(7):eabc6093.

93. Wu X, Sosunov AA, Lado W, Teoh JJ, Ham A, Li H, et al. Synaptic hyperexcitability of cytomegalic pyramidal neurons contributes to epileptogenesis in tuberous sclerosis complex. Cell Reports. 2022 Jul;40(3):111085.

94. Kostovic I, Judas M. Transient patterns of cortical lamination during prenatal life: Do they have implications for treatment? Neuroscience & Biobehavioral Reviews. 2007;31(8):1157–68.

95. Soumiya H, Fukumitsu H, Furukawa S. Prenatal immune challenge compromises development of upper-layer but not deeper-layer neurons of the mouse cerebral cortex. J of Neuroscience Research. 2011 Sep;89(9):1342–50.

96. Ikezu S, Yeh H, Delpech JC, Woodbury ME, Van Enoo AA, Ruan Z, et al. Inhibition of colony stimulating factor 1 receptor corrects maternal inflammation-induced microglial and synaptic dysfunction and behavioral abnormalities. Mol Psychiatry. 2021 Jun;26(6):1808–31.

97. Donovan APA, Basson MA. The neuroanatomy of autism - a developmental perspective. J Anat. 2017 Jan;230(1):4–15.

98. Miyamoto A, Wake H, Ishikawa AW, Eto K, Shibata K, Murakoshi H, et al. Microglia contact induces synapse formation in developing somatosensory cortex. Nat Commun. 2016 Nov;7(1):12540.

99. Thion MS, Mosser CA, Férézou I, Grisel P, Baptista S, Low D, et al. Biphasic Impact of Prenatal Inflammation and Macrophage Depletion on the Wiring of Neocortical Inhibitory Circuits. Cell Reports. 2019 Jul;28(5):1119–1126.e4.

100. Silvin A, Qian J, Ginhoux F. Brain macrophage development, diversity and dysregulation in health and disease. Cell Mol Immunol. 2023 Jun 26;10.1038/s41423-023-01053–6.

