## Supplementary material for "Influenza A virus during pregnancy disrupts maternal intestinal immunity and fetal cortical development in a dose- and time-dependent manner": MIA Reporting Guidelines

### Maternal Immune Activation Model Reporting Guidelines Checklist

| ARRIVE Reporting Guideline & Recommendation | Arrive Item | MIA Model Specific Reporting Recommendation<br>***Please complete this chart for each point outlined below. If not applicable, write N/A*** |
|---|---|---|
| **Study design**<br>➢Overview of immune activation issues<br><br>For each experiment, give brief details of the study design including:<br>a. The number of experimental and control groups.<br>b. Any steps taken to minimize the effects of subjective bias when allocating animals to treatment (e.g. randomization procedure) and when assessing results (e.g. if done, describe who was blinded and when).<br>c. The experimental unit (e.g. a single animal, group or cage of animals).<br><br>A time-line diagram or flow chart can be useful to illustrate how complex study designs were carried out. | 6 | MIA Specific Reporting:<br>a. General need for improved reporting in MIA model methods + reporting pilot data<br>○ Details on pilot data:<br>We have already published all relevant data establishing this model, which includes a Reporting Guidelines Checklist describing previous pilot data (see doi: 10.1016/j.bbi.2021.05.004). |
| **Experimental procedures**<br>➢Compounds<br>➢Validation measures<br><br>For each experiment and each experimental group, including controls, provide precise details of all procedures carried out. For example:<br>a. How (e.g. drug formulation and dose, site and route of administration, anaesthesia and analgesia used [including monitoring], surgical procedure, method of euthanasia). Provide details of any specialist equipment used, including supplier(s).<br>b. When (e.g. time of day).<br>c. Where (e.g. home cage, laboratory, water maze).<br>d. Why (e.g. rationale for choice of specific anaesthetic, route of administration, drug dose used). | 7 | Provide details of:<br>a. Compounds – source, vehicle, preparation/storage, administration route, volume administered, whether anesthetics were used at time of immune challenge.<br>○ Name of compound: Mouse-adapted influenza A virus (H3N2) strain X31<br>○ Catalogue number: N/A<br>○ Lot number: N/A<br>○ Vehicle control used: Sterile PBS; pH = 7.4<br>○ Route of administration: Intranasal<br>○ Volume administered: 1 000 or 10 000 TCID50 in 50ul<br>○ Storage conditions: Viral stock kept at -80C<br>○ Anesthetic (type, dose, duration) used:<br>Inhalation of isoflurane (gas 2.5%, 1 min)<br><br>b. Housing variables at injection - temperature of room at injection time, cage change at time of injection or not<br>○ Light cycle of animal housing room: 12 h; 7 am on, 7 pm off<br>○ Time of day of injection: Morning<br>○ Room temperature at injection time: Ambient<br>○ Did a cage change occur at time of injection: YES |

| | | |
|---|---|---|
| | | c. Validation of immune activation – behavior, physiological indices and/or cytokine data, including pilot dosing data<br>○ Method used to verify immune activation:<br>Immune activation via IAV infection in pregnant mice was validated by failure to gain weight, presence of viral expression in the lungs (qPCR), lung lesion scores (histopathology), and elevation in circulating inflammatory cytokines (Inflammatory LEGENDplex).<br><br>d. Validation of gestational timing – vaginal plug, estrous cycle, weight gain<br>○ Method of validating gestational timing:<br>Estrous cycle was initiated by exposing females to soiled male bedding. The male was introduced to the female the evening of the second day. Vaginal plugs were used to determine successful copulation and were designated as gestational day (GD)0.5. Weight gain of 2 g or more by GD9.5 (inoculation) was used to confirm successful pregnancy.<br><br>Additional comments:<br>Presence of a vaginal plug was assessed each morning for 2-to-4 consecutive days after introduction of the male stud. Following breeding, female pairs were moved into a clean cage. |
| **Experimental animals**<br>➢Species/strain/vendor<br><br>a. Provide details of the animals used, including species, strain, sex, developmental stage (e.g. mean or median age plus age range) and weight (e.g. mean or median weight plus weight range).<br>b. Provide further relevant information such as the source of animals, international strain nomenclature, genetic modification status (e.g. knock-out or transgenic), genotype, health/immune status, drug or test naïve, previous procedures, etc. | 8 | Provide details of:<br>a. Species – considerations for appropriate species (mouse, rat, non human primate, other)<br>○ Species: Mouse<br><br>b. Strain – variability in strain can influence model<br>○ Strain: C57BL/6NTac<br><br>c. Maternal/Offspring Physiological Variables at time of immune challenge – age, body weight<br>○ Maternal Age at challenge: 12-29 weeks<br>○ Maternal Body weight: 18-25 g<br>○ Offspring Age at challenge: Embryonic day (E) 9.5<br>○ Offspring Sex: Both males and females tested<br>○ Offspring Body weight: N/A<br><br>d. Vendor – even within the same strain, vendor can influence endpoints<br>○ Vendor: Taconic Biosciences<br>○ Location of Vendor: Cambridge City, IN, USA<br>○ Room/area where animals originated from: N/A |

| | | |
|---|---|---|
| | | Additional Comments:<br>All mice are murine pathogen free (MPF) status per Taconic Biosciences health standards. All females were nulliparous virgins prior to breeding. |
| **Housing and husbandry**<br>➢Cage, ventilation, bedding, enrichment<br><br>Provide details of:<br>a. Housing (type of facility e.g. specific pathogen free [SPF]; type of cage or housing; bedding material; number of cage companions; tank shape and material etc. for fish).<br>b. Husbandry conditions (e.g. breeding program, light/dark cycle, temperature, quality of water etc for fish, type of food, access to food and water, environmental enrichment).<br>c. Welfare-related assessments and interventions that were carried out prior to, during, or after the experiment. | 9 | Provide details of:<br>a. Caging systems<br>○ *At breeding*<br>Material of cage: Individually ventilated microisolator polysulfone<br>Cage dimensions: 15" L x 5" W x 7.5" H<br>○ *After parturition*<br>Material of cage: N/A<br>Cage dimensions: N/A<br>○ *At weaning*<br>Material of cage: N/A<br>Cage dimensions: N/A<br><br>b. Animal Holding room<br>○ Temperature in room: Ambient<br>○ Humidity in room: 45%<br>○ Ventilation system: Positive pressure<br>○ Specific pathogen free [SPF]: YES<br>○ Are males & females housed in the same or separate rooms:<br>Housed in separate rooms after weaning<br>c. Bedding exchanges/bedding type<br>○ Type of cage bedding used: Corncob<br>○ Frequency of cage changes per week<br>*during gestation:* Once every 2 weeks<br>*during neonatal period:* N/A<br>*following weaning:* N/A<br><br>d. Breeding - bred on site or timed pregnant, how many different sires (are the same fathers breeding with both experimental and control dams)<br>Breeding location: Bred on site. Sires were bred to experimental control dams. |

- Gestational age at shipping: N/A
- Biological age of dams (if not listed in Section 8c): see 8c
- Number of Dams bred: 225
- How many times have dams been mated previously: 0-3
- How many times did the dams mate and not become pregnant: 0-3
- Are the dams primiparous or multiparous? All dams are primiparous
- What was the frequency of maternal handling during the gestational/neonatal period (e.g. cage cleanings, weighing, blood collection manipulations): Daily handling for weighing only.
- Biological age of sires: 10-44 weeks
- Number of sires bred: 72
- How many times have sires been mated previously: 0-3
- How many times did the sires mate successfully (e.g. mating resulted in pregnancy, full term birth): 1-4
- If bred previously, what was the interval between mating times:
- Are sires matched to experimental and control dams: YES
- Describe the mating design (1:1, 1:2 etc): 1:2

e. Social enrichment – number of cage companions
   - Number of cage companions prior to breeding: Pair-housed
   - Gestational age when dam separated for parturition: GD9.5
   - Number of cage companions at weaning: N/A

f. Physical enrichment – describe enrichment devices, and when enrichment is in the cage (removed when pups born? Or present throughout study), does the enrichment type change? How frequently?
   - Describe what type of enrichment devices (and how many) are included in cage/housing room:

Pair-housed females were given a cotton nestlet to assemble a nest with. Nesting material was present pre- and post-breeding. At GD9.5, dams were inoculated and singly-housed.

Singly-housed females were moved to conventional cages and given a cotton nestlet. IAV-infected dams were provided a petri-dish containing moist chow starting at 2 days post infection (dpi).

| | | |
|---|---|---|
| | | ○ Does enrichment type/access change across study? NO<br>○ If so, when does enrichment type/access change (e.g. enrichment removed prior to parturition and replaced in late neonatal period):<br><br>Additional Comments: |
| **Sample size**<br>➢ Litter versus offspring<br><br>a. Specify the total number of animals used in each experiment, and the number of animals in each experimental group.<br>b. Explain how the number of animals was arrived at. Provide details of any sample size calculation used.<br>c. Indicate the number of independent replications of each experiment, if relevant. | 10 | Provide details of:<br>a. Maternal N vs offspring N<br>○ What is the total number of dams/litters included in the study: 100<br>○ What is the total number of offspring per litter included the study: 1 representative fetus per litter.<br>b. Litter size and sex distribution<br>○ What size was each litter maintained at: N/A<br>○ What age did culling take place at: N/A<br>○ How many males and females were maintained in each litter: N/A<br>c. Cross fostering<br>○ Did cross fostering occur: NO<br>○ If so, at what age did cross fostering occur: N/A<br><br>Additional Comments: |

| | | |
|---|---|---|
| **Allocating animals to experimental groups**<br><br>a. Give full details of how animals were allocated to experimental groups, including randomization or matching if done.<br>b. Describe the order in which the animals in the different experimental groups were treated and assessed. | 11 | a. How many offspring per litter were used in each measure: 1<br><br>b. Randomization/Matching procedures<br>○ What procedures were used to assign animals to groups:<br>Pregnant dams were randomly assigned to experimental groups (saline, IAV-X31 1 000 TCID50, or IAV-X31 10 000 TCID50) on GD9.5. One fetus per dam was chosen randomly for downstream processing.<br><br>c. Sex as a biological variable (behavioral and physiological outcomes)<br>○ Were both males and females evaluated in each behavioral and physiological outcome: YES<br><br>Additional Comments:<br>Fetal sex was not identified. However, because fetuses were randomly chosen at each endpoint, both male and female fetuses were assumed to be included in each measure. |
| **Experimental outcomes**<br>➢Behavioral testing<br>➢Physiological endpoints<br><br>Clearly define the primary and secondary experimental outcomes assessed (e.g. cell death, molecular markers, behavioral changes). | 12 | a. Maternal behavior and pup interactions<br>○ If maternal care was evaluated, were there differences following immunogen challenge (if so, please briefly describe):<br>N/A<br><br>b. Age(s) of offspring at behavioral testing/physiological evaluation endpoints:<br>N/A<br><br>c. Order of testing (e.g. behavioral test order)<br>○ Were animals evaluated in a counter-balanced order in terms of:<br>*presentation of tests to each animal:* NO<br>*order of experimental/control groups run through each test:* NO<br>○ What was the inter-test interval if a single animal underwent a battery of tests:<br>N/A |

| | | |
|---|---|---|
| | | Additional Comments: |
| **Statistical methods**<br><br>a. Provide details of the statistical methods used for each analysis.<br>b. Specify the unit of analysis for each dataset (e.g. single animal, group of animals, single neuron).<br>c. Describe any methods used to assess whether the data met the assumptions of the statistical approach. | 13 | a. Unit of analysis for each data set<br>○ Is the unit (n) of each analysis based on number of litters, or number of animals used per group:<br>Dam/litter was the experimental unit for all outcomes. |
| **Other Disclosures** | | Please make note of any other extraneous variables that you would like to report (e.g. fire alarms, construction, temporary relocations, other variables that you think we should be considering in our studies etc.): |

The recommended use of this reporting form is to fill it out and include it as supplemental material for each of your laboratory's research publications. If there are difficulties utilizing/adapting this fillable form, please contact one of the corresponding authors to request a copy. The authors give permission for this table to be edited for use in reporting on other animal models (e.g. postnatal immune challenge models, early life stress models) as appropriate.

Kentner AC, Bilbo AD, Brown AS, Hsiao EY, McAllister AK, Meyer U, Pearce BD, Pletnikov MV, Yolken RH, Bauman MD. (2018). Maternal immune activation: reporting guidelines to improve the rigor, reproducibility, and transparency of the model. Neuropsychopharmacology, https://doi.org/10.1038/s41386-018-0185-7.
