## Supplemental Figures for "Influenza A virus during pregnancy disrupts maternal intestinal immunity and fetal cortical development in a dose- and time-dependent manner"

### Slide 1
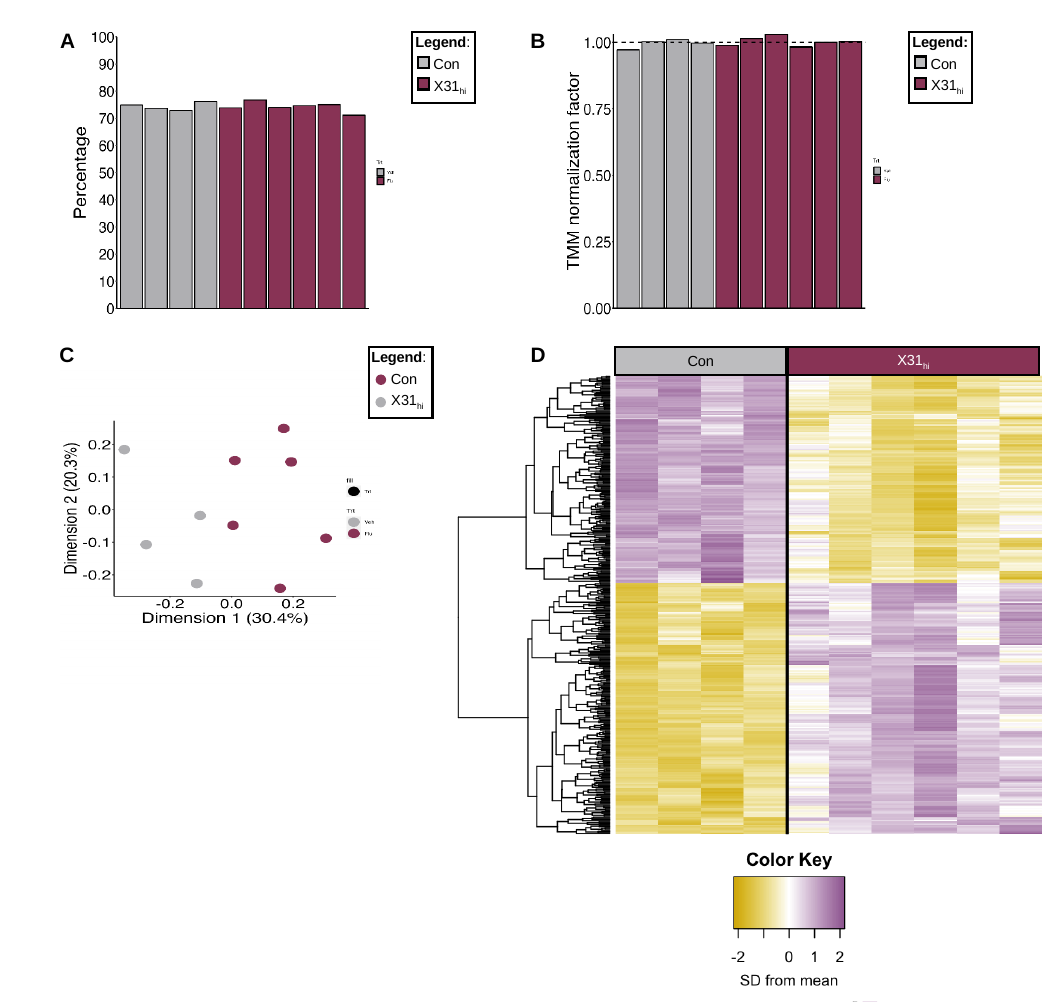

B
A
Legend:
Con
X31_hi_
Legend:
Con
X31_hi_
D
C
Legend:
Con
X31_hi_
Con
X31_hi_

### Slide 2
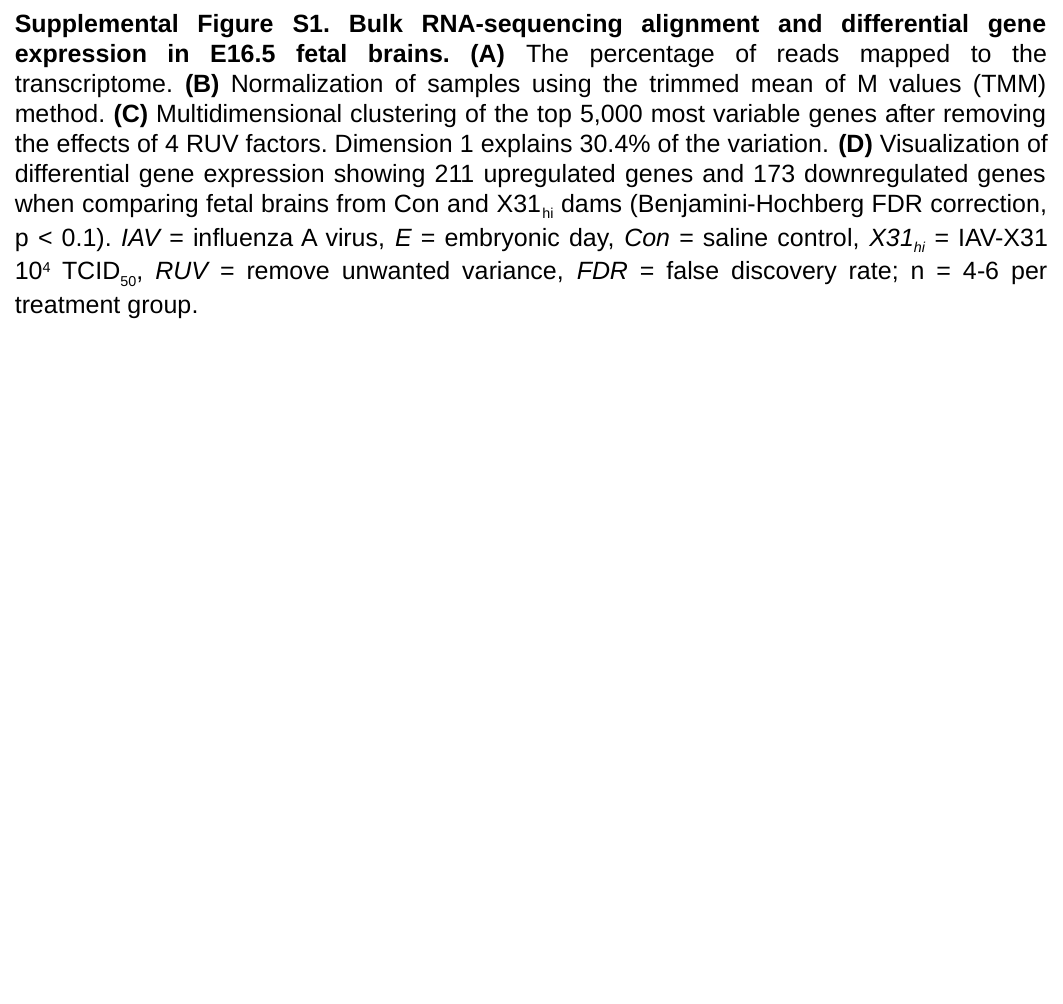

Supplemental Figure S1. Bulk RNA-sequencing alignment and differential gene expression in E16.5 fetal brains. (A) The percentage of reads mapped to the transcriptome. (B) Normalization of samples using the trimmed mean of M values (TMM) method. (C) Multidimensional clustering of the top 5,000 most variable genes after removing the effects of 4 RUV factors. Dimension 1 explains 30.4% of the variation. (D) Visualization of differential gene expression showing 211 upregulated genes and 173 downregulated genes when comparing fetal brains from Con and X31_hi_ dams (Benjamini-Hochberg FDR correction, p < 0.1). IAV = influenza A virus, E = embryonic day, Con = saline control, X31_hi_ = IAV-X31 10^4^ TCID_50_, RUV = remove unwanted variance, FDR = false discovery rate; n = 4-6 per treatment group.

### Slide 3
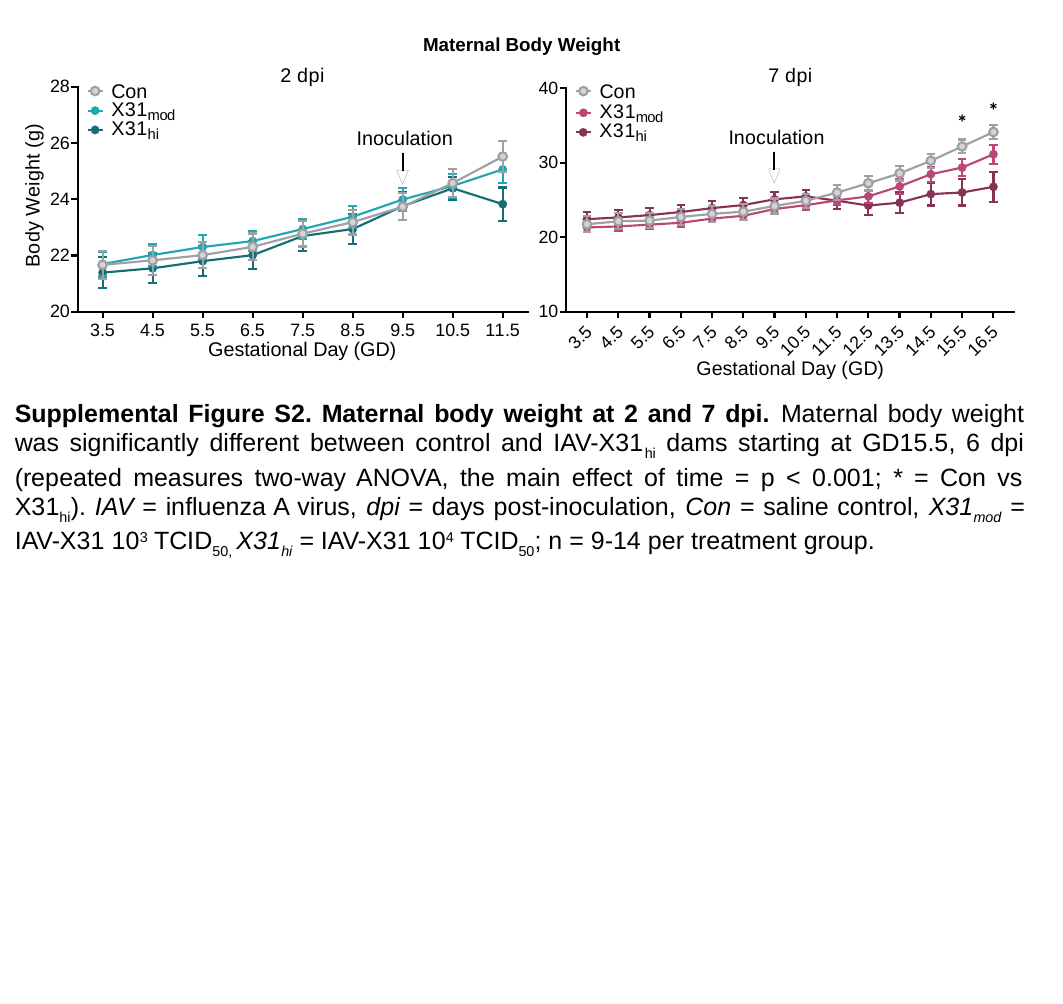

Maternal Body Weight
Supplemental Figure S2. Maternal body weight at 2 and 7 dpi. Maternal body weight was significantly different between control and IAV-X31_hi_ dams starting at GD15.5, 6 dpi (repeated measures two-way ANOVA, the main effect of time = p < 0.001; * = Con vs X31_hi_). IAV = influenza A virus, dpi = days post-inoculation, Con = saline control, X31_mod_ = IAV-X31 10^3^ TCID_50,_ X31_hi_ = IAV-X31 10^4^ TCID_50_; n = 9-14 per treatment group.

### Slide 4
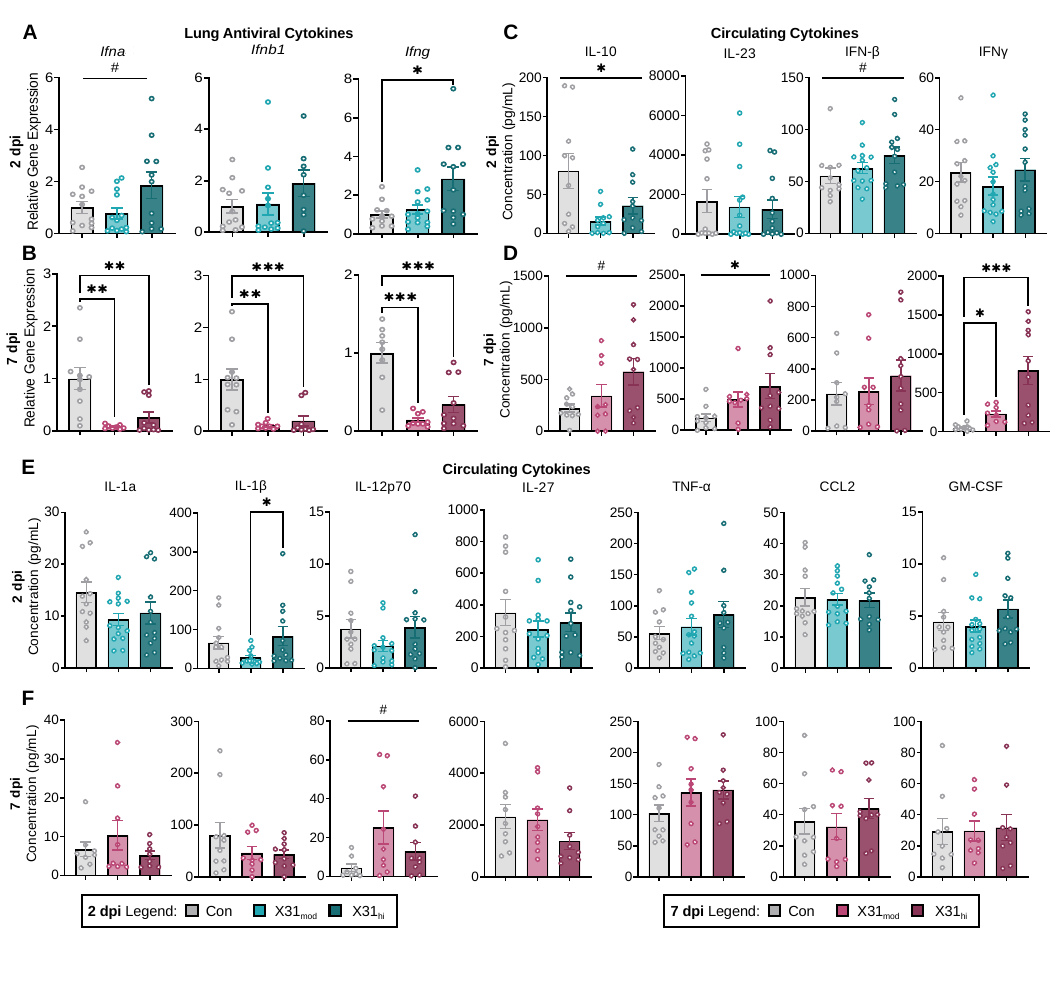

C
A
Lung Antiviral Cytokines
Circulating Cytokines
D
B
E
Circulating Cytokines
F
X31_mod_
Con
X31_hi_
7 dpi Legend:
X31_mod_
Con
X31_hi_
2 dpi Legend:

### Slide 5
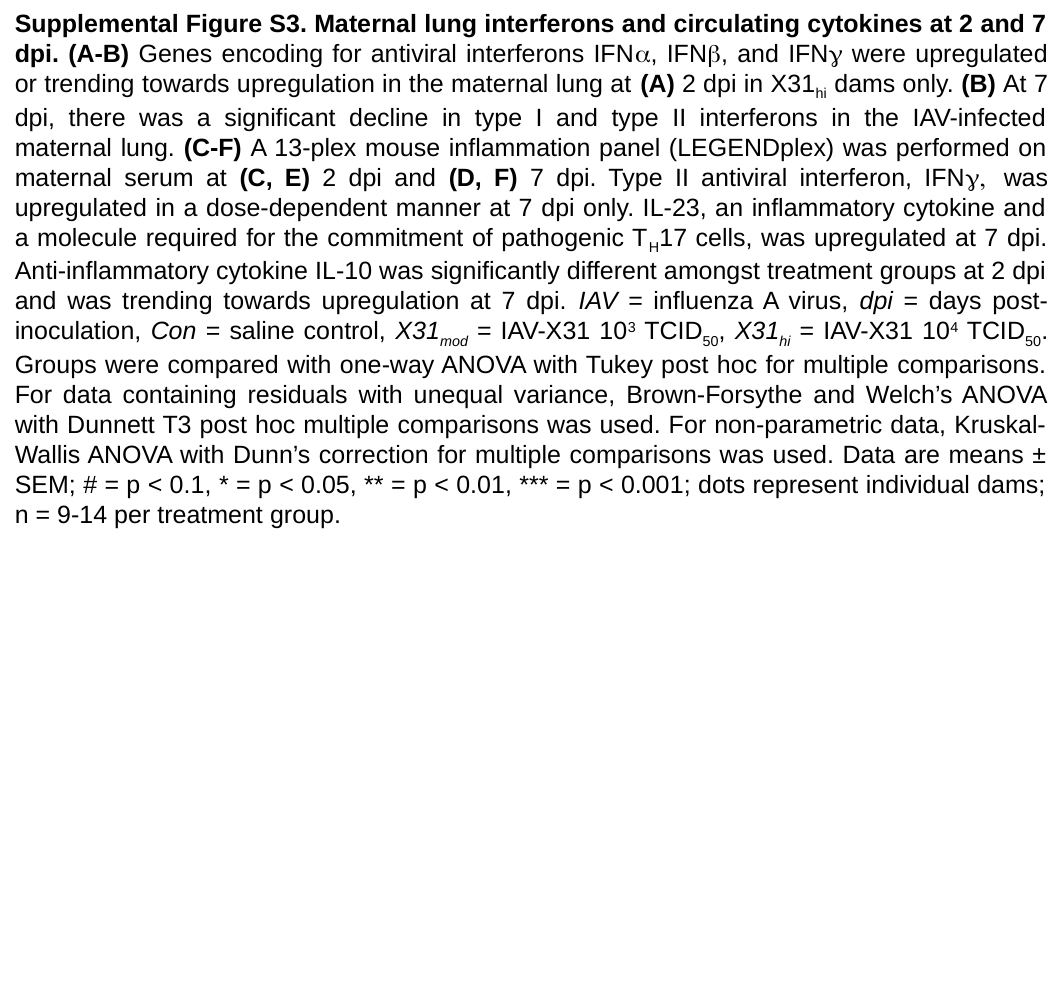

Supplemental Figure S3. Maternal lung interferons and circulating cytokines at 2 and 7 dpi. (A-B) Genes encoding for antiviral interferons IFNα, IFNβ, and IFNγ were upregulated or trending towards upregulation in the maternal lung at (A) 2 dpi in X31_hi_ dams only. (B) At 7 dpi, there was a significant decline in type I and type II interferons in the IAV-infected maternal lung. (C-F) A 13-plex mouse inflammation panel (LEGENDplex) was performed on maternal serum at (C, E) 2 dpi and (D, F) 7 dpi. Type II antiviral interferon, IFNγ, was upregulated in a dose-dependent manner at 7 dpi only. IL-23, an inflammatory cytokine and a molecule required for the commitment of pathogenic T_H_17 cells, was upregulated at 7 dpi. Anti-inflammatory cytokine IL-10 was significantly different amongst treatment groups at 2 dpi and was trending towards upregulation at 7 dpi. IAV = influenza A virus, dpi = days post-inoculation, Con = saline control, X31_mod_ = IAV-X31 10^3^ TCID_50_, X31_hi_ = IAV-X31 10^4^ TCID_50_. Groups were compared with one-way ANOVA with Tukey post hoc for multiple comparisons. For data containing residuals with unequal variance, Brown-Forsythe and Welch’s ANOVA with Dunnett T3 post hoc multiple comparisons was used. For non-parametric data, Kruskal-Wallis ANOVA with Dunn’s correction for multiple comparisons was used. Data are means ± SEM; # = p < 0.1, * = p < 0.05, ** = p < 0.01, *** = p < 0.001; dots represent individual dams; n = 9-14 per treatment group.

### Slide 6
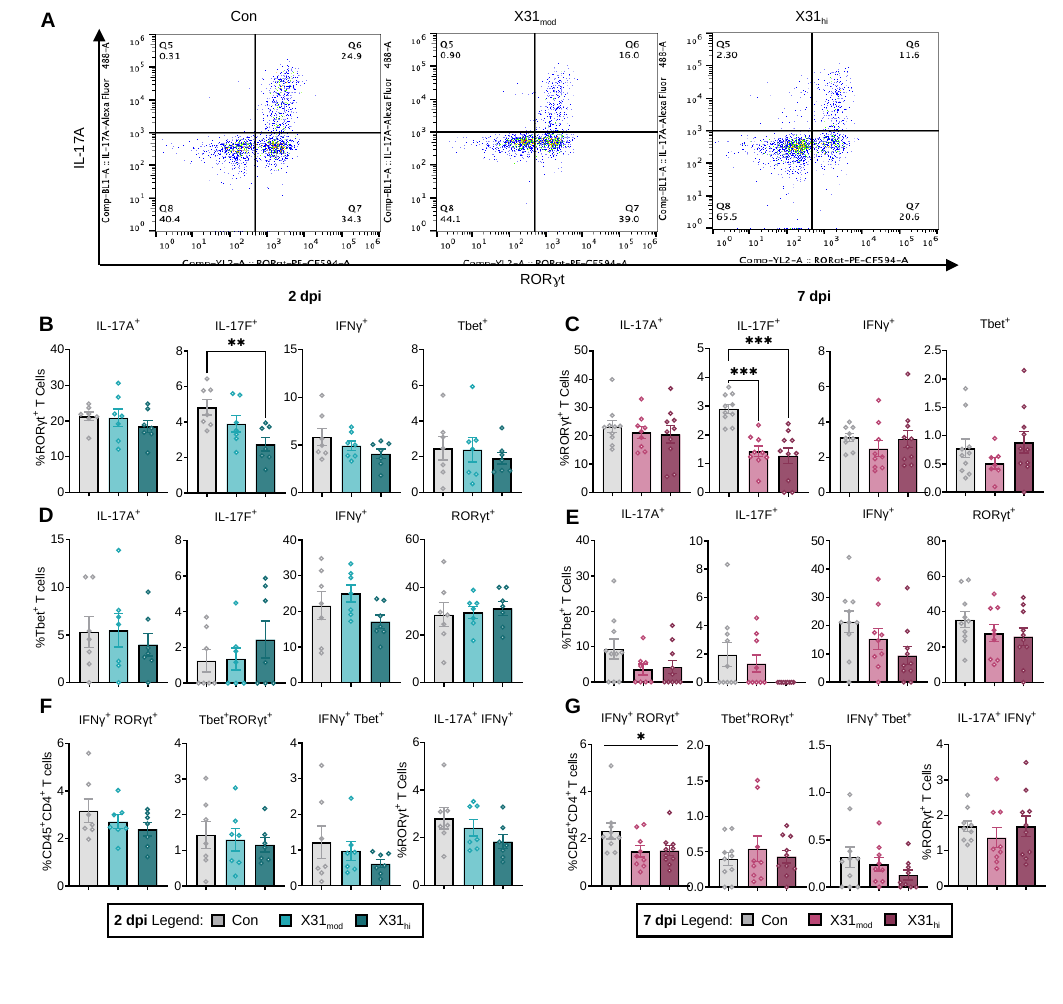

IL-17A
RORγt
X31_hi_
X31_mod_
Con
A
7 dpi
2 dpi
B
C
D
E
F
G
X31_mod_
Con
X31_hi_
7 dpi Legend:
X31_mod_
Con
X31_hi_
2 dpi Legend:

### Slide 7
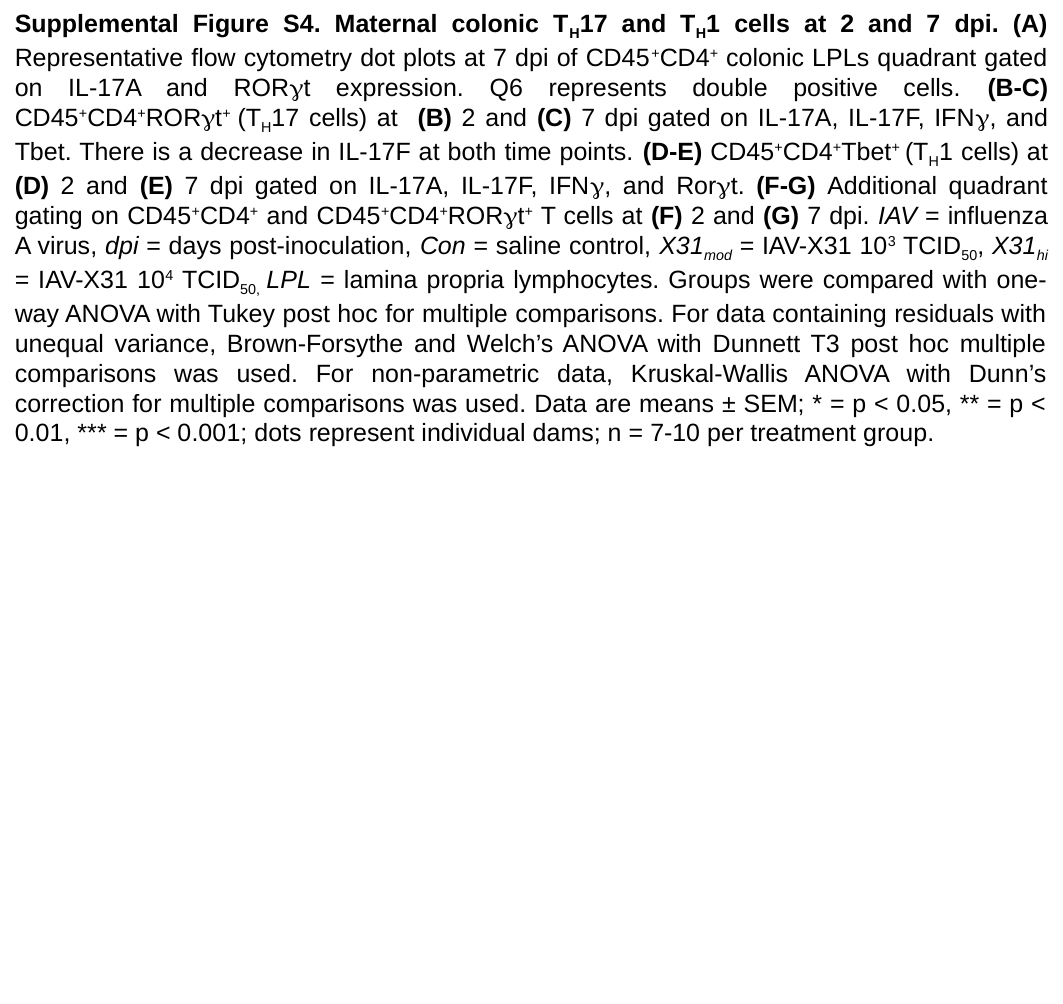

Supplemental Figure S4. Maternal colonic T_H_17 and T_H_1 cells at 2 and 7 dpi. (A) Representative flow cytometry dot plots at 7 dpi of CD45^+^CD4^+^ colonic LPLs quadrant gated on IL-17A and RORγt expression. Q6 represents double positive cells. (B-C) CD45^+^CD4^+^RORγt^+^ (T_H_17 cells) at (B) 2 and (C) 7 dpi gated on IL-17A, IL-17F, IFNγ, and Tbet. There is a decrease in IL-17F at both time points. (D-E) CD45^+^CD4^+^Tbet^+^ (T_H_1 cells) at (D) 2 and (E) 7 dpi gated on IL-17A, IL-17F, IFNγ, and Rorγt. (F-G) Additional quadrant gating on CD45^+^CD4^+^ and CD45^+^CD4^+^RORγt^+^ T cells at (F) 2 and (G) 7 dpi. IAV = influenza A virus, dpi = days post-inoculation, Con = saline control, X31_mod_ = IAV-X31 10^3^ TCID_50_, X31_hi_ = IAV-X31 10^4^ TCID_50,_ LPL = lamina propria lymphocytes. Groups were compared with one-way ANOVA with Tukey post hoc for multiple comparisons. For data containing residuals with unequal variance, Brown-Forsythe and Welch’s ANOVA with Dunnett T3 post hoc multiple comparisons was used. For non-parametric data, Kruskal-Wallis ANOVA with Dunn’s correction for multiple comparisons was used. Data are means ± SEM; * = p < 0.05, ** = p < 0.01, *** = p < 0.001; dots represent individual dams; n = 7-10 per treatment group.

### Slide 8
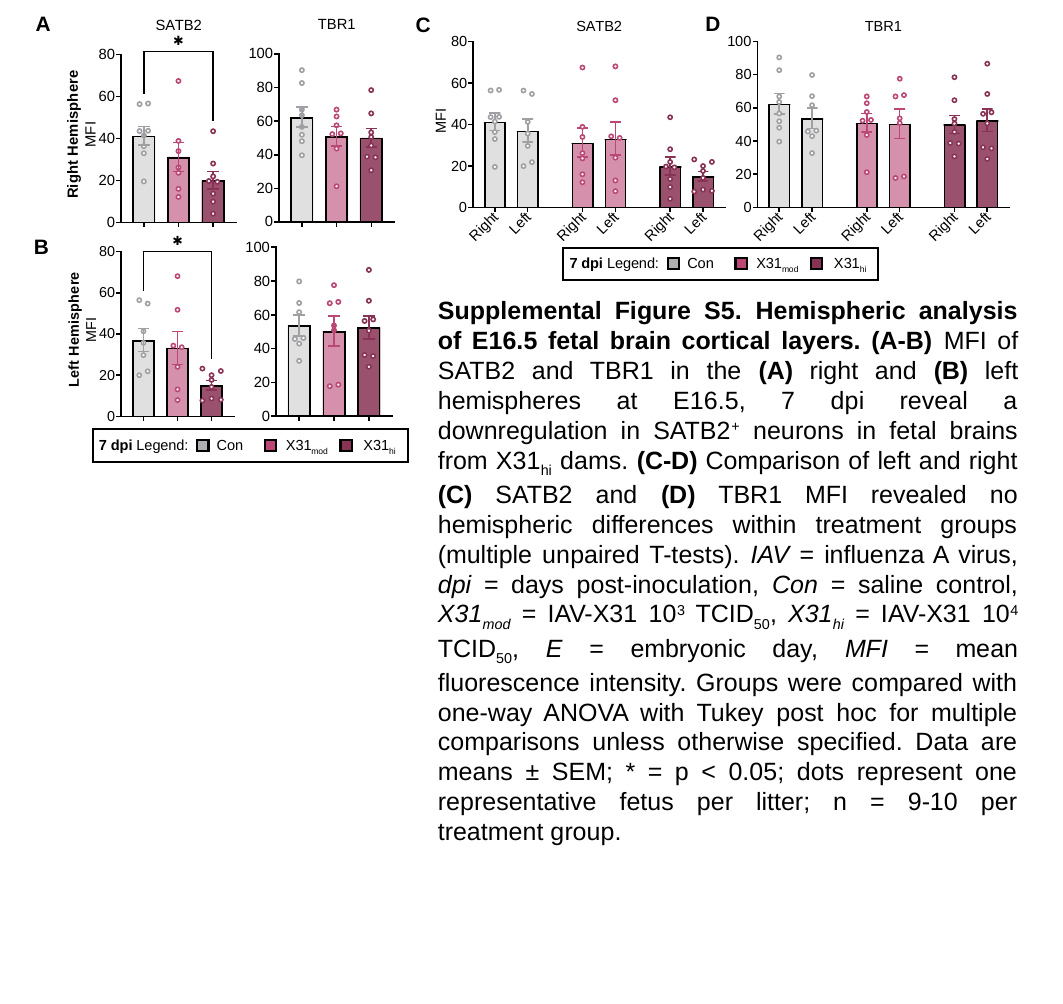

A
D
C
B
X31_mod_
Con
X31_hi_
7 dpi Legend:
Supplemental Figure S5. Hemispheric analysis of E16.5 fetal brain cortical layers. (A-B) MFI of SATB2 and TBR1 in the (A) right and (B) left hemispheres at E16.5, 7 dpi reveal a downregulation in SATB2^+^ neurons in fetal brains from X31_hi_ dams. (C-D) Comparison of left and right (C) SATB2 and (D) TBR1 MFI revealed no hemispheric differences within treatment groups (multiple unpaired T-tests). IAV = influenza A virus, dpi = days post-inoculation, Con = saline control, X31_mod_ = IAV-X31 10^3^ TCID_50_, X31_hi_ = IAV-X31 10^4^ TCID_50_, E = embryonic day, MFI = mean fluorescence intensity. Groups were compared with one-way ANOVA with Tukey post hoc for multiple comparisons unless otherwise specified. Data are means ± SEM; * = p < 0.05; dots represent one representative fetus per litter; n = 9-10 per treatment group.
X31_mod_
Con
X31_hi_
7 dpi Legend:

### Slide 9
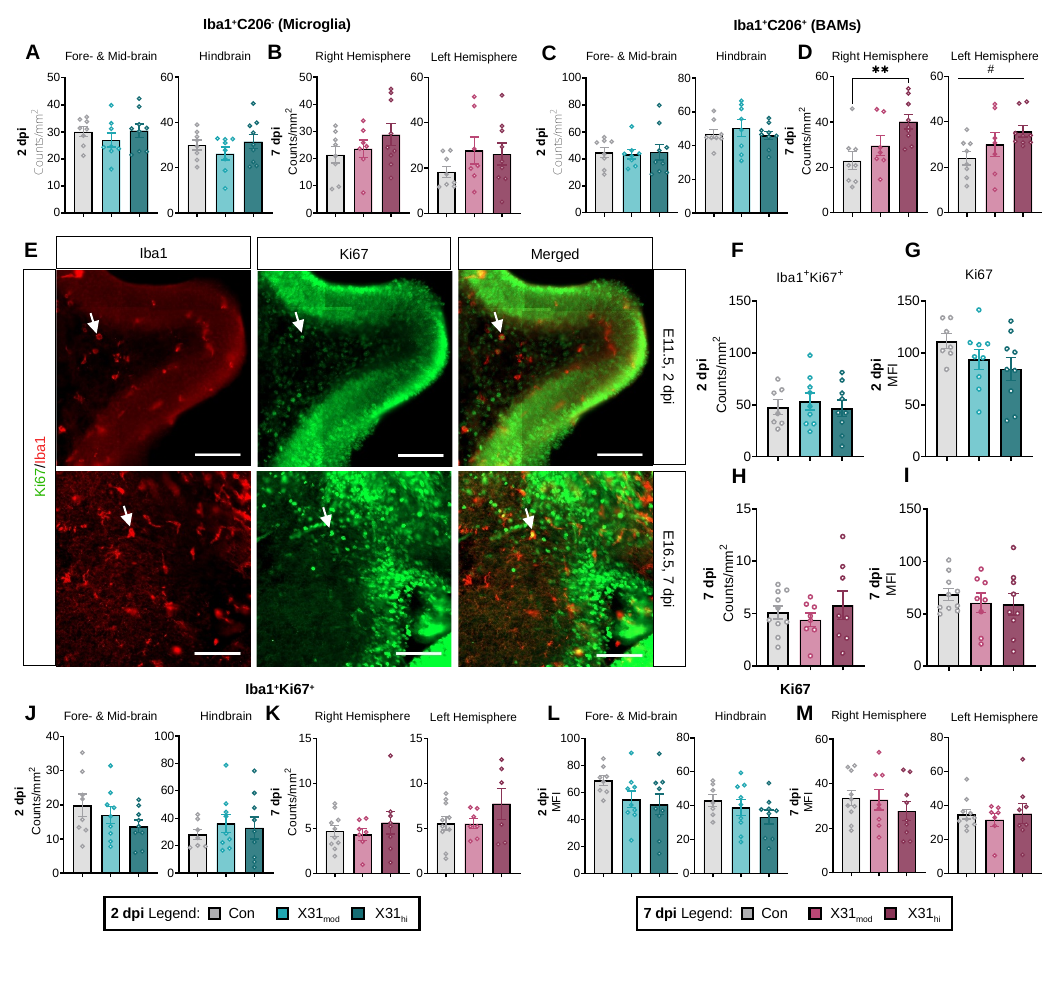

Iba1^+^C206^-^ (Microglia)
Iba1^+^C206^+^ (BAMs)
D
A
B
C
G
E
Iba1
Ki67
Merged
E11.5, 2 dpi
Ki67/Iba1
E16.5, 7 dpi
F
I
H
Iba1^+^Ki67^+^
Ki67
J
K
M
L
X31_mod_
Con
X31_hi_
7 dpi Legend:
X31_mod_
Con
X31_hi_
2 dpi Legend:

### Slide 10
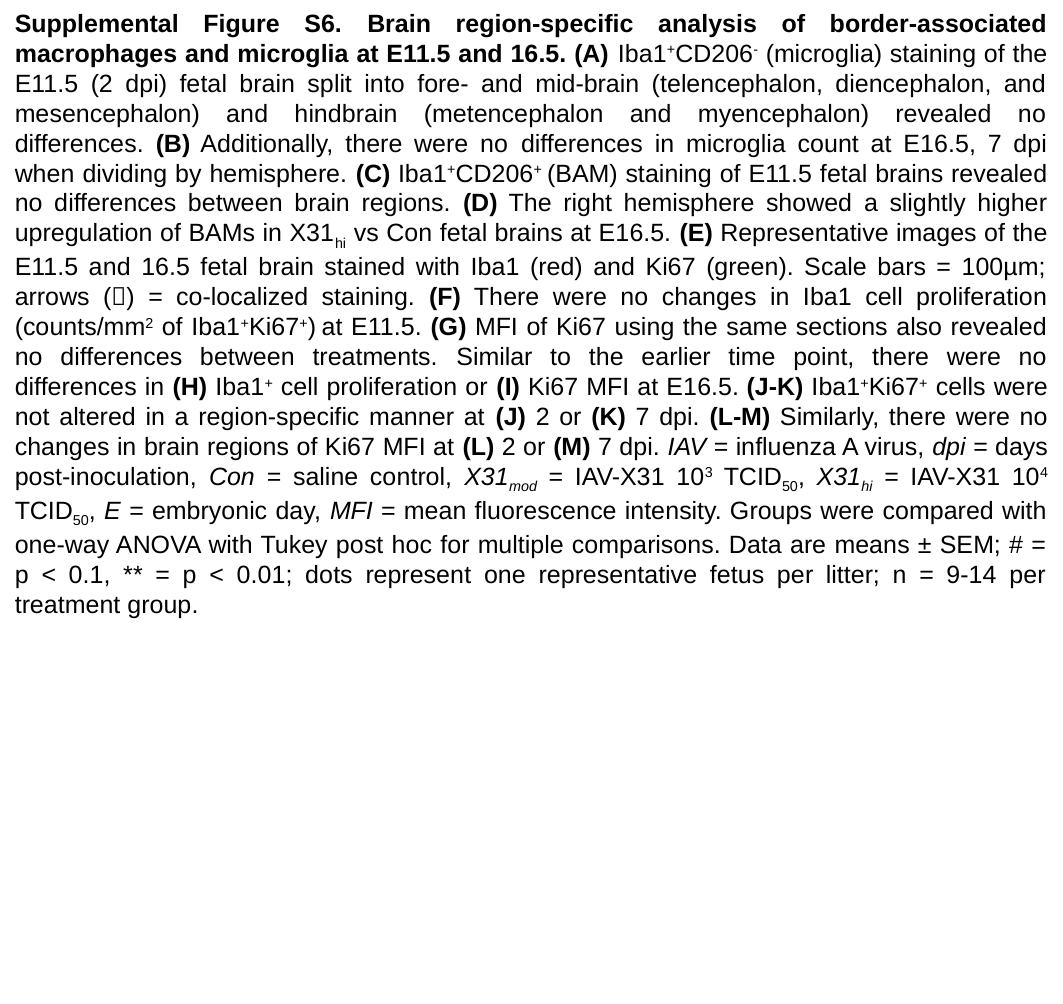

Supplemental Figure S6. Brain region-specific analysis of border-associated macrophages and microglia at E11.5 and 16.5. (A) Iba1^+^CD206^-^ (microglia) staining of the E11.5 (2 dpi) fetal brain split into fore- and mid-brain (telencephalon, diencephalon, and mesencephalon) and hindbrain (metencephalon and myencephalon) revealed no differences. (B) Additionally, there were no differences in microglia count at E16.5, 7 dpi when dividing by hemisphere. (C) Iba1^+^CD206^+^ (BAM) staining of E11.5 fetal brains revealed no differences between brain regions. (D) The right hemisphere showed a slightly higher upregulation of BAMs in X31_hi_ vs Con fetal brains at E16.5. (E) Representative images of the E11.5 and 16.5 fetal brain stained with Iba1 (red) and Ki67 (green). Scale bars = 100µm; arrows (🡪) = co-localized staining. (F) There were no changes in Iba1 cell proliferation (counts/mm^2^ of Iba1^+^Ki67^+^) at E11.5. (G) MFI of Ki67 using the same sections also revealed no differences between treatments. Similar to the earlier time point, there were no differences in (H) Iba1^+^ cell proliferation or (I) Ki67 MFI at E16.5. (J-K) Iba1^+^Ki67^+^ cells were not altered in a region-specific manner at (J) 2 or (K) 7 dpi. (L-M) Similarly, there were no changes in brain regions of Ki67 MFI at (L) 2 or (M) 7 dpi. IAV = influenza A virus, dpi = days post-inoculation, Con = saline control, X31_mod_ = IAV-X31 10^3^ TCID_50_, X31_hi_ = IAV-X31 10^4^ TCID_50_, E = embryonic day, MFI = mean fluorescence intensity. Groups were compared with one-way ANOVA with Tukey post hoc for multiple comparisons. Data are means ± SEM; # = p < 0.1, ** = p < 0.01; dots represent one representative fetus per litter; n = 9-14 per treatment group.
